# Mechanism of tissue expansion in the early stage of the migration

**DOI:** 10.1101/2021.07.20.453119

**Authors:** Abhimanyu Kiran, Navin Kumar, Vishwajeet Mehandia

**Author notes:** **Correspondence:** Abhimanyu Kiran & Vishwajeet Mehandia, &.

## Abstract

The collective cell migration is observed in many biological processes such as wound healing, embryogenesis, and cancer metastasis. Despite extensive theoretical and experimental studies on collective cell motion, there is no unified mechanism to explain it. In this work, we experimentally report the collectively growing cell colonies in the sub-marginal region of a freely expanding cell monolayer. These colonies could be responsible for the highly aligned collective cell migration observed in front cell rows. Our results provide a basic framework to understand the physical mechanism responsible for collective cell migration in the freely expanding monolayer.

## 1 Introduction

The mechanism involved in the single-cell migration is well known, where a cell forms a new focal adhesion at the leading end, followed by retraction, and then detachment of focal adhesion from the trailing end (Friedl and Gilmour, 2009). Its migration is affected by external parameters like substrate stiffness, topography, elasticity, and porosity (Friedl et al., 2004; Rørth, 2009). Just like single cells, the migration of monolayer is affected by its biophysical and biochemical environment. However, the collective cell migration cannot be introduced as many single cells migrating in the same direction with a constant velocity (Ilina and Friedl, 2009) because each cell is connected to its neighbour through cell-cell adhesion and actin cytoskeleton (Mayor and Etienne-Manneville, 2016). Therefore, the direction of cell migration is influenced by the neighbouring cell (Vicsek et al., 1995; Szabo et al., 2006), resulting in much more efficient migration as compared to single cells put together without considering their intercellular interactions (Mayor and Etienne-Manneville, 2016). It indicates that intercellular forces play a vital role in the unknown mechanism by which multiple cell rows communicate to achieve collective cell motion (Ladoux, 2009).

Earlier, the collective cell migration was assumed to be driven by the leader cells actively pulling their follower cells (Du Roure et al., 2005; Gov, 2007; Poujade et al., 2007; Vitorino and Meyer, 2008). Later, Xavier et al. (Trepat et al., 2009a) experimentally demonstrated the presence of high traction forces up to ~10 cell rows behind the marginal cells. These forces are heterogeneous (Serra-Picamal et al., 2012) and may cause the cells to move randomly or even in the opposite direction as that of leader cells (Petitjean et al., 2010; Vedula et al., 2012; Reffay et al., 2014). The unequal distribution between the leader and follower cells may lead to a never-ending tug of war (Trepat et al., 2009a; Tambe et al., 2011; Serra-Picamal et al., 2012). It is assumed that the vigorous leader cells dominate the tug of war and can pull the follower cells in the forward direction. Still, the amount of force (~100 nN) exerted by the leader cells is not sufficient to pull the entire monolayer (Ilina and Friedl, 2009; Reffay et al., 2014). Thus, the leader cells do require some assistance from the follower cells or some hidden cooperative mechanism by which they can communicate with the follower cells and migrate effectively as a sheet. Several studies have shown the vital role of follower cells in collective cell migration. These studies include the propagation of a slow mechanical wave through the monolayer (Serra-Picamal et al., 2012), high correlation in normal stress fluctuation (Tambe et al., 2011), and correlated velocity field up to 10-15 cell rows from the leading edge (Petitjean et al., 2010). Additionally, Farooqui and Fenteany reported the presence of ‘cryptic’ groups of cells behind the leading edge of the wound that extends their lamellipodia in the basal region of their front cells (Farooqui and Fenteany, 2005). These sub-marginal cells could be involved in the collective cell migration.

Further, the biomechanics of cells have been extensively analyzed to understand the parameters that govern the migration of epithelial cell aggregates such as an island, stripes, and complete monolayer. Zimmermann et al. employed a particle-based simulation approach to model expanding cell monolayer (Zimmermann et al., 2016). They successfully captured the force pattern developed by the cell aggregates, but their model lacks the physiological features of the real cells. Magno et al. established the link between cell surface mechanics and tissue dynamics using simulation and analytical results (Magno et al., 2015). Coburn et al. performed a computational analysis of the epithelial cell islands and stripes using the vertex model and contact inhibition of locomotion (CIL). They predicted the decrease in the basal protrusion, traction force, and apical ratio from the edge to the island centre (Coburn et al., 2016). While the junctional and principle stress increase in the same direction (Maruthamuthu et al., 2011; Jasaitis et al., 2012; Mertz et al., 2013; Coburn et al., 2016). The increase of island size leads to an increase in the junctional stress at the centre (Coburn et al., 2016) and traction force at the rim (Mertz et al., 2012). The expansion of the island would reduce its height as per the conservation of volume (Coburn et al., 2016). The biomechanical analysis of epithelial cell aggregates provides excellent insight into intercellular forces and their relation with traction forces. However, the role of the follower cells and the mechanism by which they contribute to collective cell migration is not well understood.

Here, we conducted live-cell imaging experiments at the leading site (~8-10 cell rows) of the Madin-Darby Canine Kidney (MDCK) cell monolayer to decipher the role of the follower cells in the expansion of the epithelial monolayer. We have found the groups of collectively growing cells in the sub-marginal region of the monolayer (the word ‘growth’ represents the increase in the projected cell area with time (Figure 6A)). In this work, we investigated the leading site of the freely-expanding epithelial monolayer to decipher the physical mechanism that could be responsible for the collective cell migration in freely expanding epithelial monolayer.

## 2 Materials and Methods

### 2.1. Cell culture and microscopy

The live-cell time-lapse experiments were carried out using MDCK *II* cells stably transfected with Green Fluorescent Protein E-Cadherin (Nelson Lab.). The cells were cultured in the growth medium made of 90% Dulbecco’s Modified Eagle Medium (DMEM) containing 10% Fetal Bovine Serum (FBS) and antibiotics. The monolayer was formed by seeding the suspended cells in the centre of a 35 mm diameter glass-bottom Petri dish (Eppendorf®). After the formation of the confluent monolayer, the growth medium was replaced with the image acquisition medium made of 90% L-15, 10% FBS, and antibiotics.

For observation, an inverted microscope (Leica DMI 6000B) integrated with a full cage incubator was employed to maintain the temperature of 37°C. The experiments were performed by keeping the height of the z-stack from 2-3 μm in such a way that the best focus of the monolayer lies at the centre of the stack. It prevents the monolayer from getting out of the focus during the experimental observation. The images were automatically captured after the interval of 5 minutes up to 4 hours using Leica’s LAS X software. After obtaining the z-stack images, the z-stack images were merged into one image using the average intensity projection at each time step.

### 2.2. Identification of growing cell colonies

The images of the experiment were skeletonized (Figure S1). Then the random numbers were assigned to the cells for their identification in a reference image at t = 0 hr (Figure 1A). The changes in the area and the geometric centre were manually measured for all cells using the ‘Wand tool’ of ImageJ. The cells that were not skeletonized properly; their change in area was manually measured by using the ‘Polygon section tool’ of ImageJ. All the measurements (change in the area and geometric centre of each cell) of the experiments were stored in an excel sheet. These measurements were used for all the analyses.

**Figure 1.**
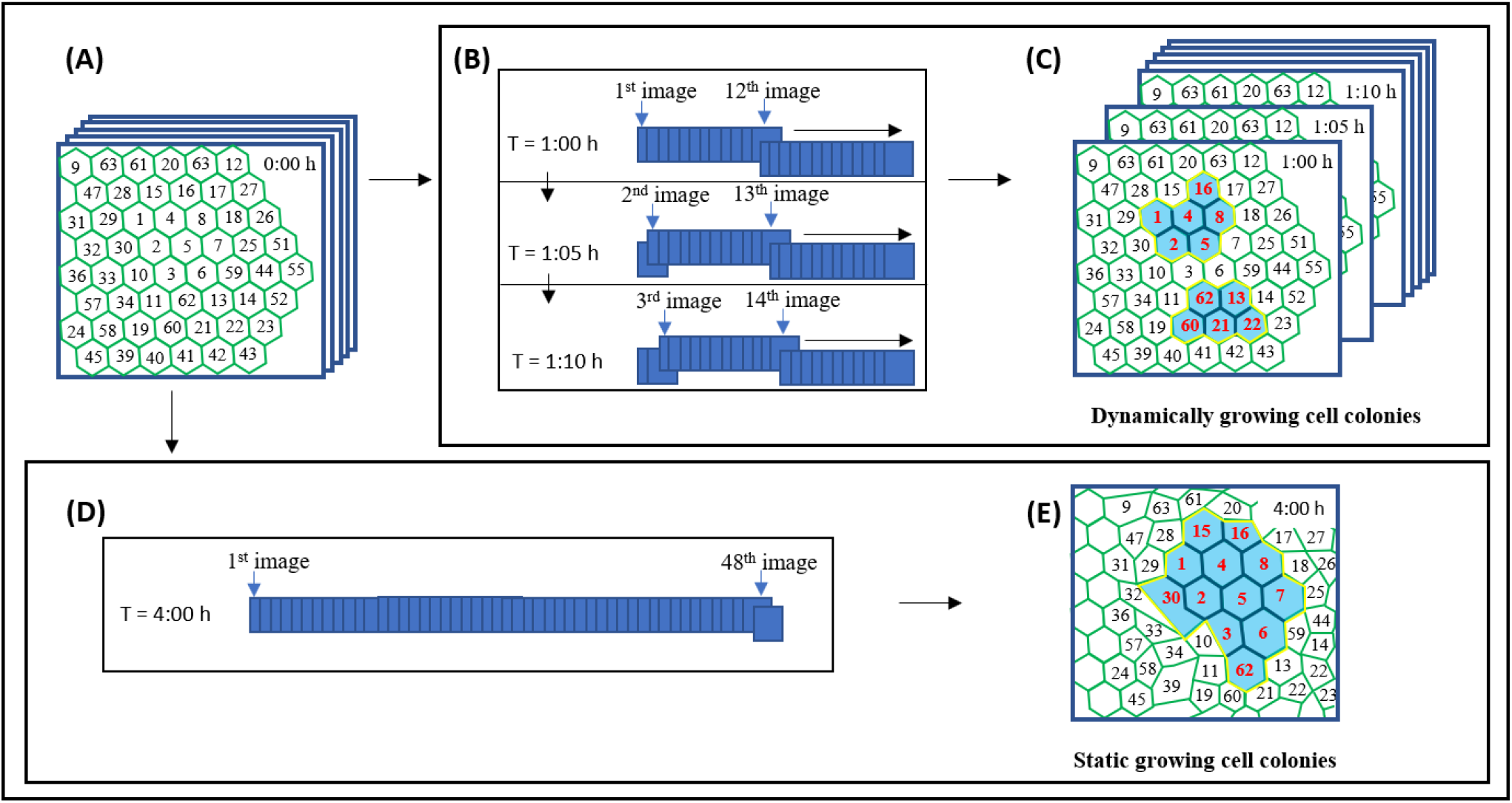
Identification of expanding colonies (A) Assigning random numbering to the cells, (B) one-hour (or 12 images) moving window analysis, (C) identification of dynamic colonies throughout the image sequence, (D) four-hour window analysis, (E) identification of the static growing colony at t = 4 hr. Note: In (C) and (E), the growing cells marked with red color, common junction of growing cells exhibiting strong Pearson correlation marked by black lines, and colonies shown by blue color region.

#### 2.2.1. Dynamically growing cell colonies

The cells expand or grow at different rates and for different durations. In order to capture these dynamics, a one-hour moving window analysis was performed where previous 12 (or one-hour) images were compared to identify the collectively growing cells. For this analysis, two eligibility criteria were considered. First was the selection of a growth criterion for growing cells. It is reported in the literature that cells of the MDCK monolayer exhibit ± 20% area fluctuation in 4 hours (Zehnder et al., 2015). Therefore, any cell that grows more than 20% of its initial area in 4 hours could be considered as a growing cell (Kiran et al., 2021). Accordingly, for one-hour moving window analysis, any cell that grows more than 5% of its initial area in one hour was marked as a growing cell (Kiran et al., 2021). The second criteria was to determine the strong positive Pearson correlation (*r_ab_* > 0.7) between the change in the area of the connected cells with time. The Pearson correlation coefficient (*r_ab_*) was measured by using the formula given below.

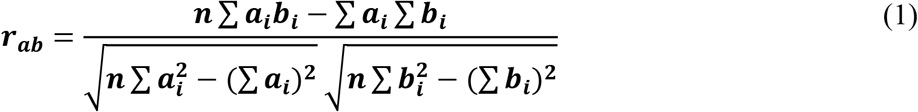

Where *r_ab_* represents the Pearson’s correlation coefficient between cell *a,* and cell *b*, *n* indicates the number of time frames or images or window size (n = 12, 24, or 36), *a_i_* and *b_i_* denotes the area of the cell *a*, and cell *b* at the *i^th^* frame, respectively.

Thereby, combining both the criteria, all the growing cells (5% growth criteria) exhibiting strong Pearson correlation (r_ab_ > 0.7) among themselves were grouped together to represent the colonies. Minimum three cells were required to form a colony.

In the moving window analysis (Figure 1B), for t = 1 hr, 1^st^ to 12^th^ images were considered, and all the growing cells exhibiting correlated growth were enclosed to form the growing colonies in the 13^th^ image (Figure 1C). Further, the moving window slide in time by dropping the 1^st^ image and picking the 13^th^ image (Figure 1B). Then, for t = 1:05 hr, 2^nd^ to 13^th^ images were considered to form the growing colonies in the 14^th^ image (Figure 1C). Similarly, this fixed sliding window step by step move up to 4hr. After the growing colonies were identified for all the image sequences, it was found that different shapes of colonies emerged at each time step (Figure 2, and Movie S1). This represents the dynamic nature of these colonies. Therefore, these colonies were named as Dynamically growing cell colonies. Since the moving window analysis requires the previous 12 (or one-hour) images, the dynamic growing colonies cannot be obtained for the first 12 images. The shape of the colonies in the moving window analysis depends on the number of previous images taken (12, 24, or 36) to identify the colonies in the current image (Figure S2).

**Figure 2.**
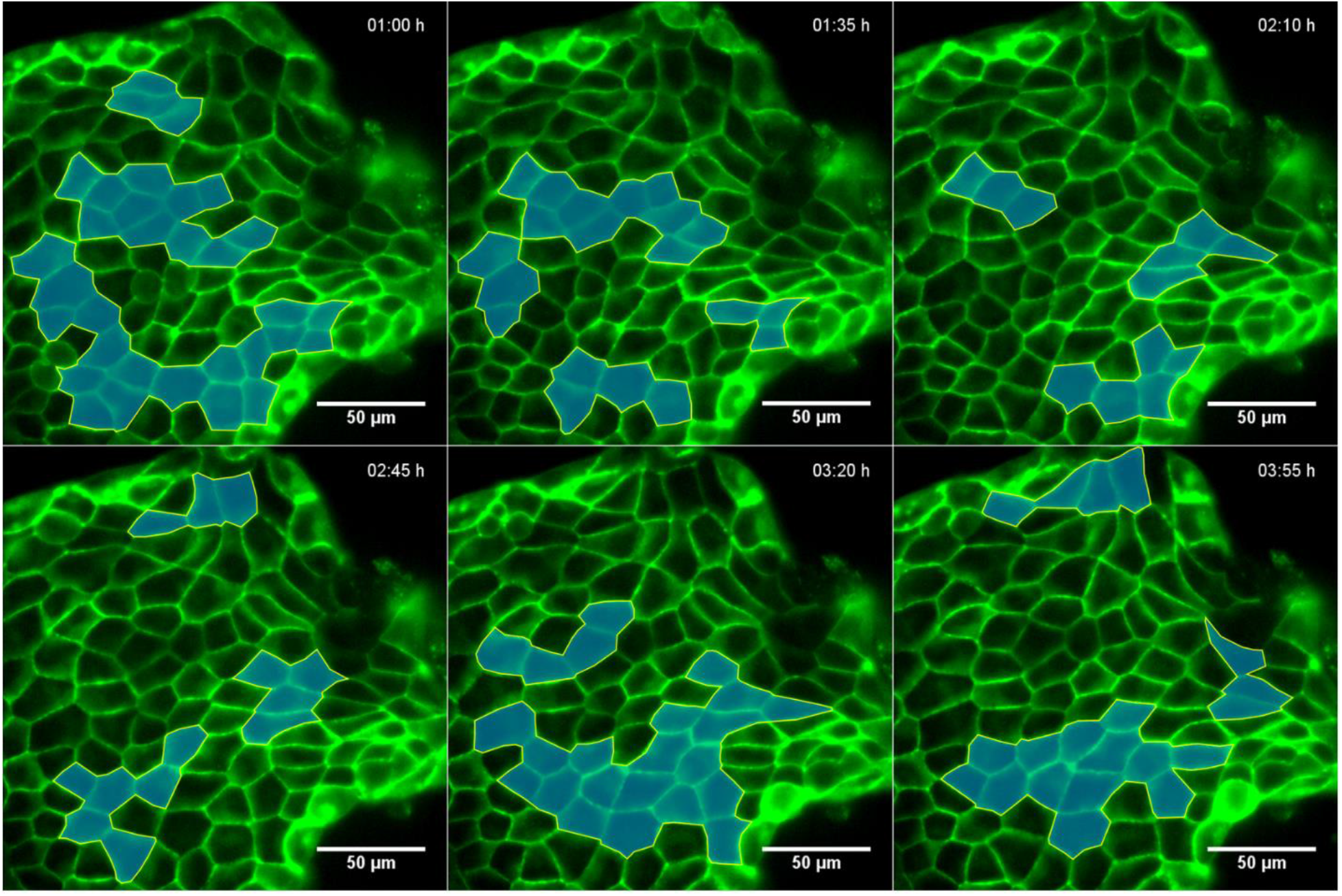
Dynamically growing cell colonies at the leading site of the freely expanding monolayer. Note: The cells of the growing colonies are represented in blue.

#### 2.2.2. Static growing cell colonies

Since it was difficult to analyze the Dynamically growing colonies (that can vary in shape, size, and location with time), therefore, for the ease of analysis, the dynamic cell colonies were converted into static cell colonies that have fixed shapes. Thereby, the largest window size (48 images or four-hour) that can cover the entire experimental span and cannot slide further was selected (Figure 1D). In this four-hour window, all the cells that grow more than 20% of their original area (criteria for growing cell) and exhibit strong Pearson correlation (*r_ab_* > 0.7) between them were grouped to form a fixed shape colony in the 49^th^ image and referred to as Static growing colonies (Figure 1E). It contained the cells which grew together in their area throughout the image sequence. It was observed that a few first row cells become leaders, which pull their immediate follower cells present in the second row. Therefore, cells of the first two rows were not considered as part of these colonies. The Dynamic and Static growing colonies were found in five independent experiments. The cells undergone division were not considered during the formation of colonies. Interestingly, bigger Static cell colonies (Figure 3A) were observed by considering a larger time span (48 images or 4 hr) as compared to Dynamic cell colonies (Figure S2) obtained at a shorter time span (12, 24 or 36 images). It may happen because the cells of the monolayer fluctuate in the short time span but grow in the long time span. Therefore, larger windows unable to capture the dynamic nature of the growing cell colonies.

**Figure 3.**
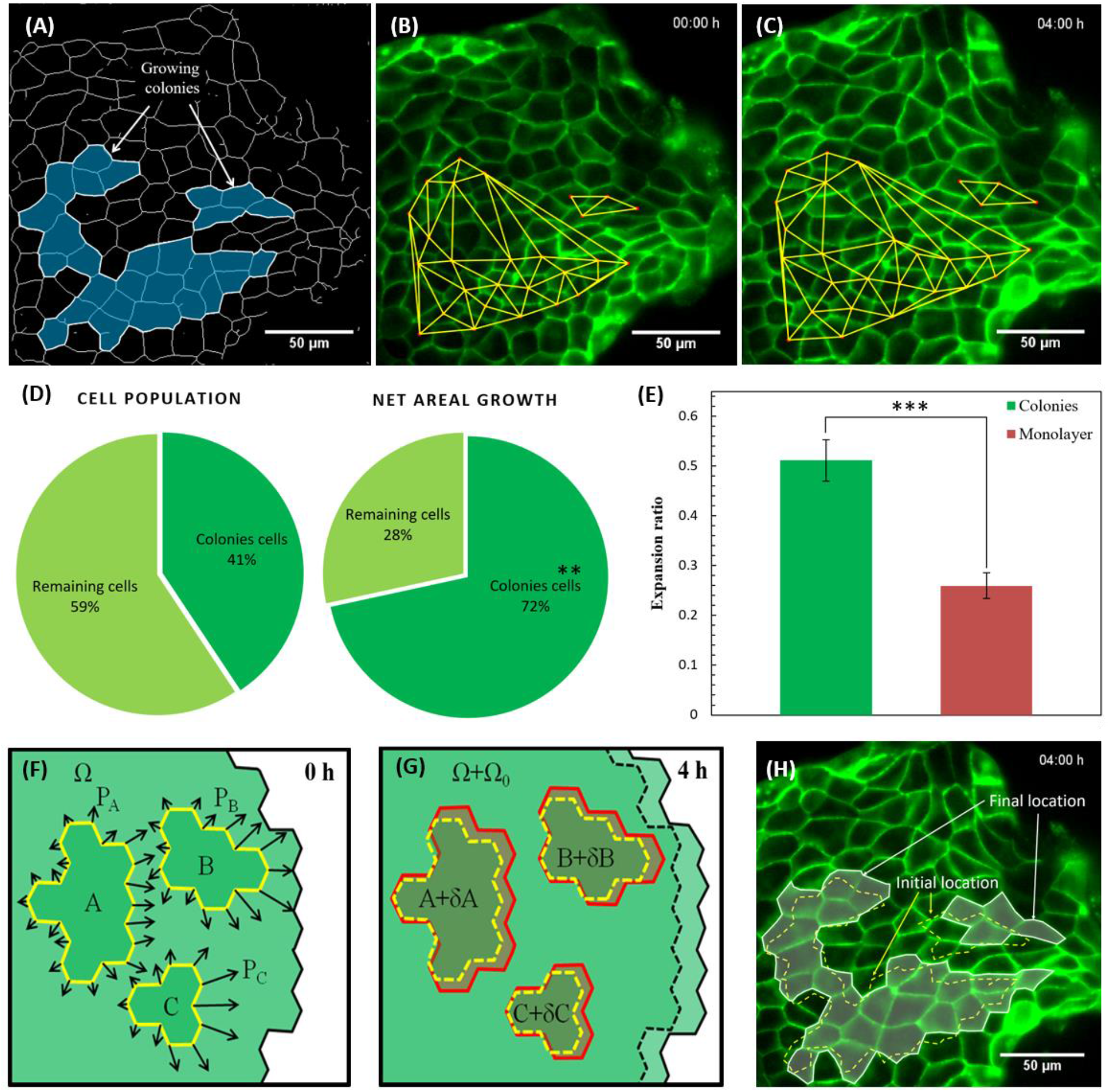
Growing colonies at the leading site (A) The growing colonies are represented in a skeletonized image, (B-C) Delaunay triangulation algorithm. The increase in the size of wire mesh from (B) t = 0 hr to (C) t = 4 hr indicates the areal expansion of the colonies, (D) cell population, and net areal growth distribution for the expanding colonies and remaining cells, (E) the expansion ratio of colonies and the monolayer (colonies and remaining cells combined), (F) the expanding colonies A, B, and C exerts intercellular forces P_A_, P_B_, and P_C_ to its neighboring cells, respectively, (G) colonies expand in the area (A+*δ*A, B+*δ*B, C+*δ*C), and (H) comparison between the final and initial location of the colonies. The dotted yellow and solid white line represents the initial and final location of the expanding colonies, respectively. Note: Five independent experiments were analyzed to obtain the result shown in (D) and (E). The images showed in (A), (B), (C), and (H) are taken as a representative experiment out of five independent experiments.

### 2.3. MATLAB codes and Algorithms

The details of the geometric centre of each cell with time were imported in a self-written MATLAB® algorithm to plot Delaunay’s triangulation and the velocity field. Similarly, the details of the change in the area of each cell with time were imported in other self-written MATLAB® algorithms to compute the average strain, growth rate, and displacement rate of various rows of the epithelial monolayer. A similar analysis was performed for five independent experiments.

### 2.4. Statistical analysis

The statistical analysis was performed using paired Student’s t-test and repeated ANOVA as suitable. The p-values were used to determine the significant differences, where *, **, and *** denotes the significance level of p < 0.05, p < 0.01, and p < 0.001, respectively. Statistical analysis was carried out using SPSS (v.21, SPSS Inc., Chicago, IL, USA) and Microsoft Office Excel (2007).

## 3 Results

### 3.1 Growing cell colonies at the leading site of freely expanding monolayer

The dynamically growing cells colonies were identified at the leading site of freely expanding monolayer (refer to “Dynamically growing cell colonies” in the Material and Method section and Figure 1B-C). Since it was difficult to analyze these dynamic cell colonies (which frequently changed their shape, number of cells, and location), therefore, the dynamic cell colonies were converted to static cell colonies (refer to “Static growing cell colonies” in the Material and Method section, Figure 1D-E, and Figure 3A). The advantage of analyzing the Dynamically growing colonies as Static growing colonies is that it considers the same cells throughout the experiment. Each cell of the Static colonies grows at a different rate. All further analyses were performed on Static growing colonies.

The Delaunay triangulation algorithm (Figure 3B-C and Movie S2) was employed to show the overall areal growth of the colonies with time. It connects the geometric centre of the cell to its neighbouring cells by drawing a link between them. The increase in the size of the wire mesh from t = 0 hr (Figure 3B) to t = 4 hr (Figure 3C) confirmed the growth of the colony. The analysis showed that only 41% of cells of the leading site contributed to 72% in the net expansion of the leading site (Figure 3D). Whereas, remaining 59% of the cells contributed to 28% in the net expansion of the leading site (Figure 3D). This indicated that cells of these colonies were the major contributor to the net expansion of the leading site. The t-test showed that even though there was no significant difference (p = 0.1262) in the population of the colonies cells and the remaining cells, but the net areal expansion of colonies cells was statistically significant (p = 0.0015) to the remaining cells. Further, the expansion ratio was determined by change in area per unit original area. It was found that the expansion ratio of the colonies (Figure 3E) was statistically significant to the expansion ratio of the leading site (colonies and remaining cells together). It also indicates that expanding colonies were the major contributor to the overall expansion of the leading site. Therefore, we hypothesized that these expanding colonies exert a considerable amount of force on their surrounding cells (Figure 3F). We also assumed that these colonies exert maximum force on their front rows (Figure 3F) and expand in size by displacing them towards the leader cells (Figure 3G-H).

As per literature, the only faction of marginal cells becomes the leader cells. These leader cells exhibit considerable lamellipodia protrusion and show outgrowth in the wake as migration progress (Vishwakarma et al., 2018). In our experiments, marginal cells that exhibited outgrowth during their migration were considered leader cells. It was often seen that velocity vectors of the cells located in the head of colonies move towards the leader cells (Figure S3A, and Figure 4A). In contrast, the velocity vectors of cells located at the tail of colonies may move in different directions as that of leader cells (marked by orange circle in Figure 4A). Therefore, we assumed that the cells of the colonies might expand in random directions by pushing their neighbouring cells. It may displace few neighbouring cells in the opposite direction as that of leader cells (red circles in Figure 4A). Interestingly, highly aligned velocity vectors (marked by a blue ellipse in Figure S3A and Figure 4A) were seen in the front rows, and we assume that these rows are being pulled by the leader cells and pushed by the expanding colonies. It suggests that the collective cell expansion of the colonies may contribute to the highly aligned velocity field observed in front rows. Further, the overall growth was evaluated by superimposing the initial (Figure S3A) and final images (Figure S3B). By comparing the initial and final shape of the colonies, it was observed that these colonies could expand in different directions, but prefer to grow towards the leader cells (Figure S3C-D and Figure 4B). Further, the correlation between the direction of growth rate and velocity was determined. A positive correlation was obtained for colonies, and a negative correlation was obtained for surrounding tissue (Figure S4). It suggests that colonies expand and migrate in the same direction, which may cause the effective displacement of the front rows towards the leader cells. It could be the reason for highly aligned velocity vectors observed in the front rows of the growing colonies (Figure S3A and Figure 4A).

**Figure 4.**
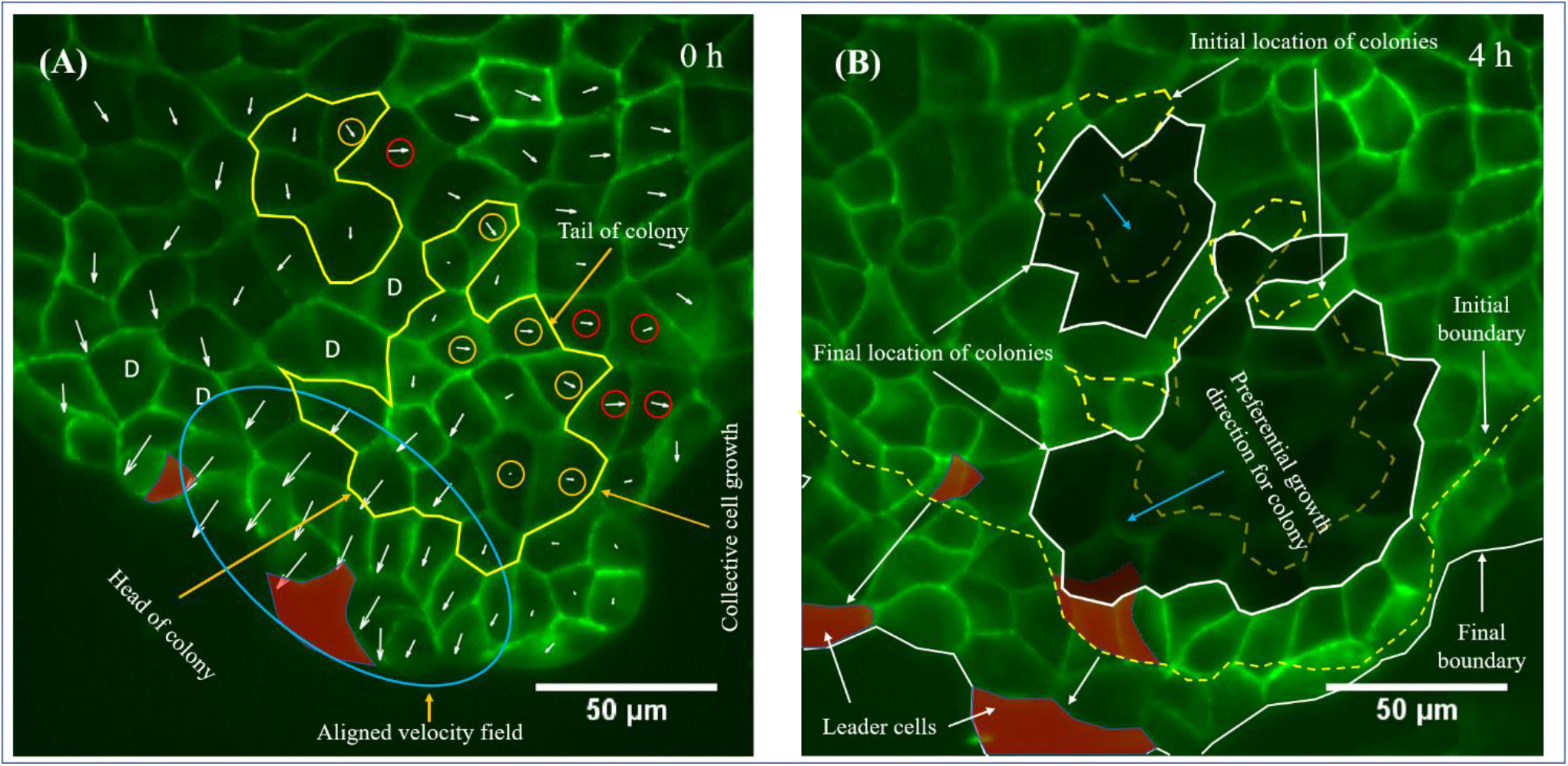
Preferential growth direction for colonies (A) Velocity field distribution at the head and tail of the colonies, and (B) preferential growth direction for the colonies. Note: Orange and red circle in (A) marks the cells of expanding colonies and neighboring cells that moves in a different direction with respect to the leader cells. The blue ellipse shows the highly aligned velocity field. The static cell colonies were analyzed here.

### 3.2 Mechanism of differential growth rate at the leading site

The leading site of monolayer migrates as a sheet, and a limited number of cell rearrangements are observed over a long-time scale. Therefore, we assumed that the cells behave like semi-solid bodies (consist of cytosol and cytoskeleton) and have some finite rigidity to shear. As discussed earlier, the expanding colonies exist among the follower cells (Figure 5A). The local growth rate of the expanding colony with area A is given by dA/dT = γ(r, t) A. The differential growth rate between the colonies and their surrounding tissue can be expressed in terms of change in pressure (Shraiman, 2005). The incremental change in pressure for the monolayer having non-uniform growth with unconstrained boundaries is proportional to the difference between local and average growth rates (Shraiman, 2005).

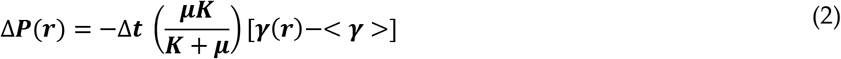

Where ΔP is the incremental change in pressure, Δt is an incremental change in time, μ is the shear rigidity, *K* is the bulk modulus, < γ > is the average growth rate of cells at the leading site, and γ(r) is the growth rate of the colony or surrounding tissue.

**Figure 5.**
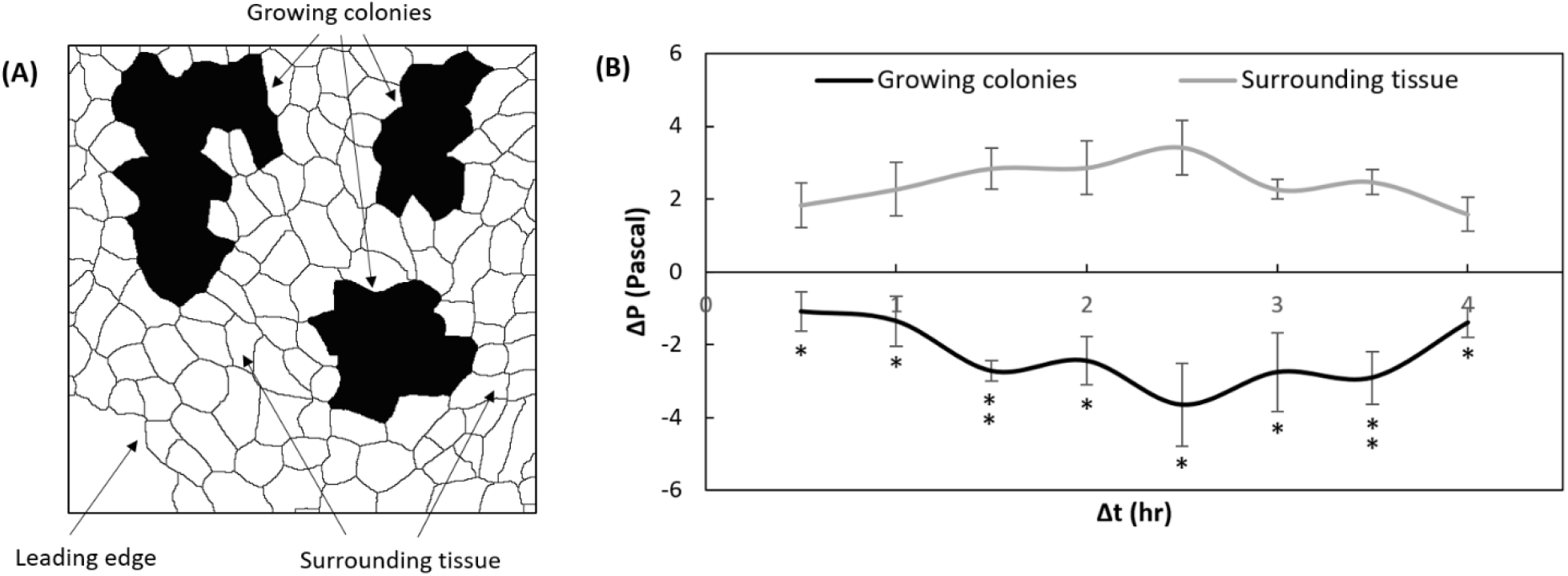
Analysis of differential growth rate at the leading site (A) Skeletonised image showing growing colonies and surrounding tissue, and (B) incremental change in the pressure for the growing colonies and surrounding tissue versus time. Note: Standard error bar plotted in (B) obtained from five independent experiments. The paired t-test was applied between the Δp values of growing colonies and surrounding tissue at the corresponding time points. These analyses were done on static cell colonies.

Taking Δt = 0.5 hr, γ(r) for expanding colonies/surrounding tissue and < γ > is determined from the experiments. The value of *μ* = 300 Pa and *K* = 200 Pa was taken, as reported in the literature (Notbohm et al., 2016). After calculating the incremental change in pressure for the colonies and surrounding tissue, it was shown as a function of time (Figure 5B). The negative ΔP obtained for the colony indicates extension, and the positive ΔP obtained for the surrounding tissue indicates compression. The incremental change in the pressure of the growing colony can be measured by Equation 3.

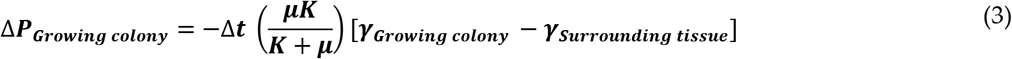

Since the cells of these colonies expand (or grow) faster than the surrounding tissue, these colonies must push their neighbouring cells. As this push originate due to differential growth rate among the follower cells. It can be termed as follower push (Equation 4).

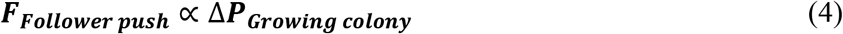

The push applied by these colonies on their neighboring cells is of compressive nature and it is reported in the literature that compressive stress (externally applied) can lead to the formation of the leader cells at the periphery of the wound margin (Janet et al., 2012). Therefore, we assume that compressive forces applied by these colonies might be responsible for recruiting the leader cells from the marginal cells. Later, these newly formed leader cells start to pull their immediate follower cells (or front cell rows) that are already being pushed by the colonies. Since pull generated by leader cells and the push from the follower cells act in the same direction. Thus, these forces might work collectively to displace the front cell rows (Equation 5).

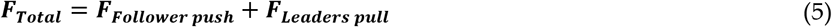

The cells of the colonies might expand to fill the space generated by the displacement of the front cell rows to maintain the tissue integrity. It may be the reason that colonies expand towards the leader cells (Figure S3C-D and Figure 4B).

### 3.3 The 3D image analysis of 2D images

The cells of the monolayer may grow, fluctuate or shrink in their area (Figure 6A). For this analysis, cells that expanded more than 20% to their initial area by the end of the observation were referred to as growing cells. Whereas the cells showed initial expansion followed by shrinkage such that the net expansion remained less than 20% of its initial area were classified as fluctuating cells. However, the cells with less final area than the initial area were considered shrinking cells. Then, the growing cells with a strong Pearson correlation (r_ab_ > 0.7) were considered part of the colony. Further, the growing cells of the colony, growing cells outside the colony, fluctuating and shrinking cells were marked in the final reference image at t = 4 hr (Figure 6B).

**Figure 6.**
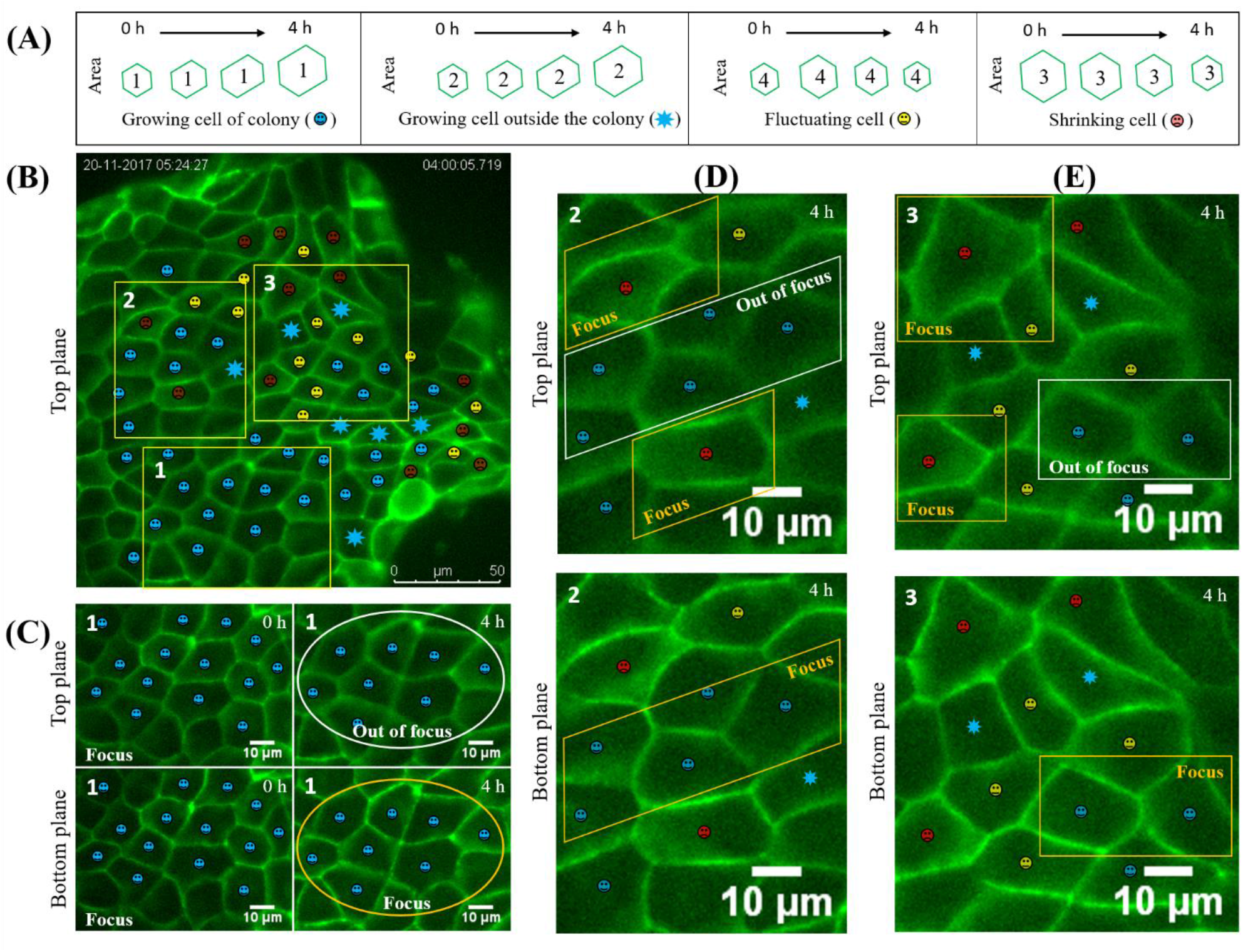
The z-stack image analysis (A) Schematic representation of growing cell of the colony, growing cell outside the colony, shrinking and fluctuating cell with time, (B) top plane image of the experiment at t = 4 hr indicating cells of the expanding colony with blue, growing cells outside the colony with the blue star, shrinking cells with red, and fluctuating cells with yellow, (C) zoom-in of window1 shown in (B), it consists of all the growing cells. At t = 0 hr, cells were focused on both the top and bottom plane. Later, at t = 4 hr, cells went out of the focus in the top plane and remained focused in the bottom plane, (D) zoom-in of window2 shown in (B), it consists of both colony cells and surrounding cells. At t = 4 hr, all cells were focused in the bottom plane, but expanding cells were out of focus, and shrinking cells remained focus in the top plane, (E) zoom-in of window3 shown in (B), it also consists of both colony cells and surrounding cells. At t = 4 hr, all cells were focused on the bottom plane, but expanding cells were out of focus on the top plane. However, shrinking cells remained focused in the top plane.

The z-stack image analysis (Figure S5) was performed to explore the 3D perspective of the 2D images. The study aimed to check the real meaning of “growth or expansion” in the colonies. For the z-stack images, the cell is focused when its apical area lies between the top and bottom plane of the z-stack. Further, few rectangular windows were created in the reference image to mark the location to be analyzed later in the z-stack (Figure 6B). The window1 consisted of all the cells of the growing colony because we want to check their change in height with time. At t = 0 hr, the cells were focused on both planes (Figure 6C). Later, at t = 4 hr, cells went out of focus from the top plane but remained focused on the bottom plane (Figure 6C). The focus and out-of-focus regions were identified by drawing the line profile across the edges and measuring the grey-scale values (Figure S6). The focused regions contained the edges that exhibited sharp peaks with high grey-scale values. However, the out-of-focus regions contained the edges that exhibited broad (or poor) peaks with low grey-scale values. Since the cells of the growing colonies became out of focus on the top plane and remained focus on the bottom plane with time. Thereby, it is assumed that cells of the colonies reduced in height with time. Since the cells of the colonies also expand in the area with time (Figure S5B). It suggests that these cells were spreading with time. The spreading of the colonies must have pushed their neighbouring cells. It may be the reason for shrinking (red colour) or fluctuation (yellow colour) in the area of their neighbouring cells with time. It can be observed by evaluating the final image of the top plane for window2 and window3, where cells of the colonies were out of focus, but neighbouring cells were focused (Figure 6D-E). The colony cells were not focused in the final image of the top plane because these cells may have decreased in their height with reference to their neighbouring cells. Further, X-Z slices (Figure S7) were also analyzed for the height, and the results were found in agreement with the results obtained from the top plane of X-Y slices (Figure 6D-E). It was observed from the X-Z slices that the common edge of the colonies was not visible on the top, while the edge of the shrinking and fluctuating cells was visible throughout the stack (Figure S7). Since the cells of the growing colonies increase in area and decrease in height, meanwhile, their immediate neighbouring cells decrease in area and increase in height (with respect to the cells of growing colonies) by the end of observation. It suggests that the spreading of the colonies might be responsible for the shrinking of their neighbouring cells.

Further, we assumed that the expansion of these sub-marginal colonies might trigger a few marginal cells to become the leaders. These leader cells exhibit considerable lamellipodia protrusion and an outgrowth. Meanwhile, these leader cells also pull the follower cells present in the 2^nd^ row. It is also reported in literature that the height of the first two cell rows is less than the average height of the monolayer (Tambe et al., 2011). Initially, at t = 0.5 hr, the leader cells were out of focus as it spread to invade the freely available space and pulled the cells of the 2^nd^ row, which were visible in both the planes (Figure 7A-B). After 1 hr, the 2^nd^ row cells become out of focus on the top plane and become poorly focused on the bottom plane (Figure 7C-D), suggesting their reduction in height. As these 2^nd^ row cells also increased in their area from t = 0.5 hr (Figure 7A-B) to t = 1.5 hr (Figure 7C-D). It suggests that the cells of 2^nd^ row cells undergone stretching as a result of the pull generated by the leader cells (Figure S5D). Here, the two major observations were (1) cells of the colonies spread to push their neighbouring cell with time, and (2) The pull generated by the leader cells cause the stretching of their immediate follower cells present in the 2^nd^ row.

**Figure 7.**
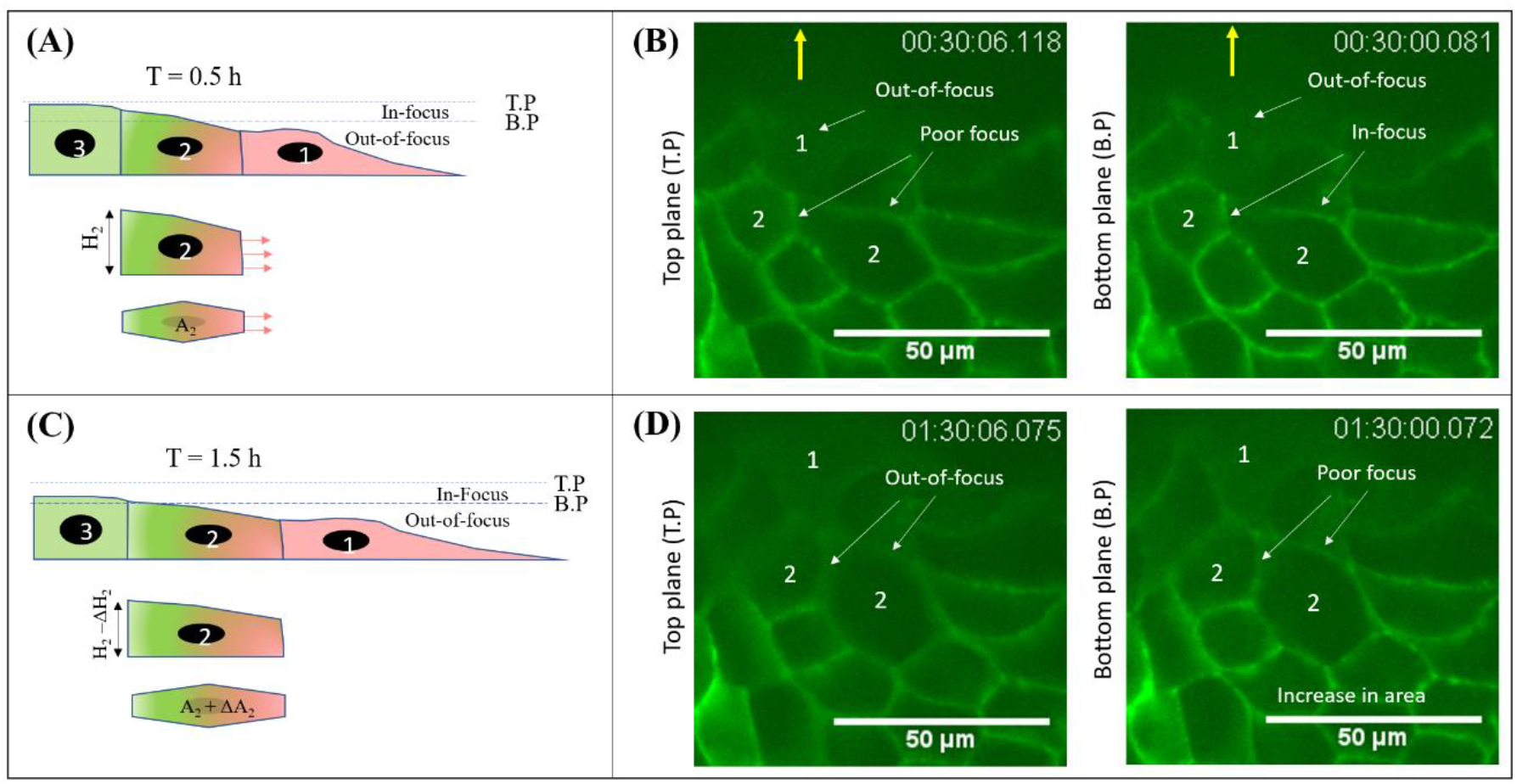
Stretching of the 2^nd^ row cells by the leader cells (A) Schematic representation of before stretching of 2^nd^ row cells, (B) top and bottom plane before stretching, (C) schematic representation of after stretching, (D) Top and bottom plane after stretching. Note: Leader cell and 2^nd^ row cells were marked by ‘1’ and ‘2’, respectively.

### 3.4 Role of followers in the migration of epithelial monolayer

The collective cell behaviour is a group phenomenon that can only be understood by analyzing all the cells together as a function of space and time. Therefore, the leading site of the monolayer was segmented into rows (Figure 8A). The average value of growth rate (GR), displacement rate (DR), and average strain (S) was computed for all cell rows. The GR is defined as the rate of change in cell area-averaged over space and time (Equation 6). The DR is defined as the rate of change of the cell’s geometric centre averaged over space and time (Equation 7). The S is defined as the change in area per unit original area-averaged over time and space (Equation 8).

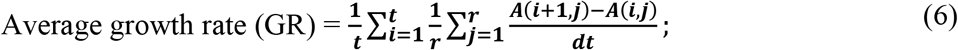

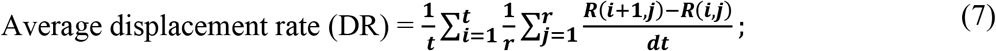

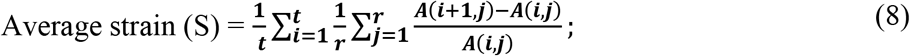

Where *A* represents the cell area, *R* is the centroid 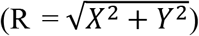, *X* & *Y* are geometric center coordinates, *t* is the number of time frames in four hours (*t = 50*), *r* is the number of cells in a row, *i* and *j* are the variables that vary from 1 to *t* and 1 to *r*, respectively.

**Figure 8.**
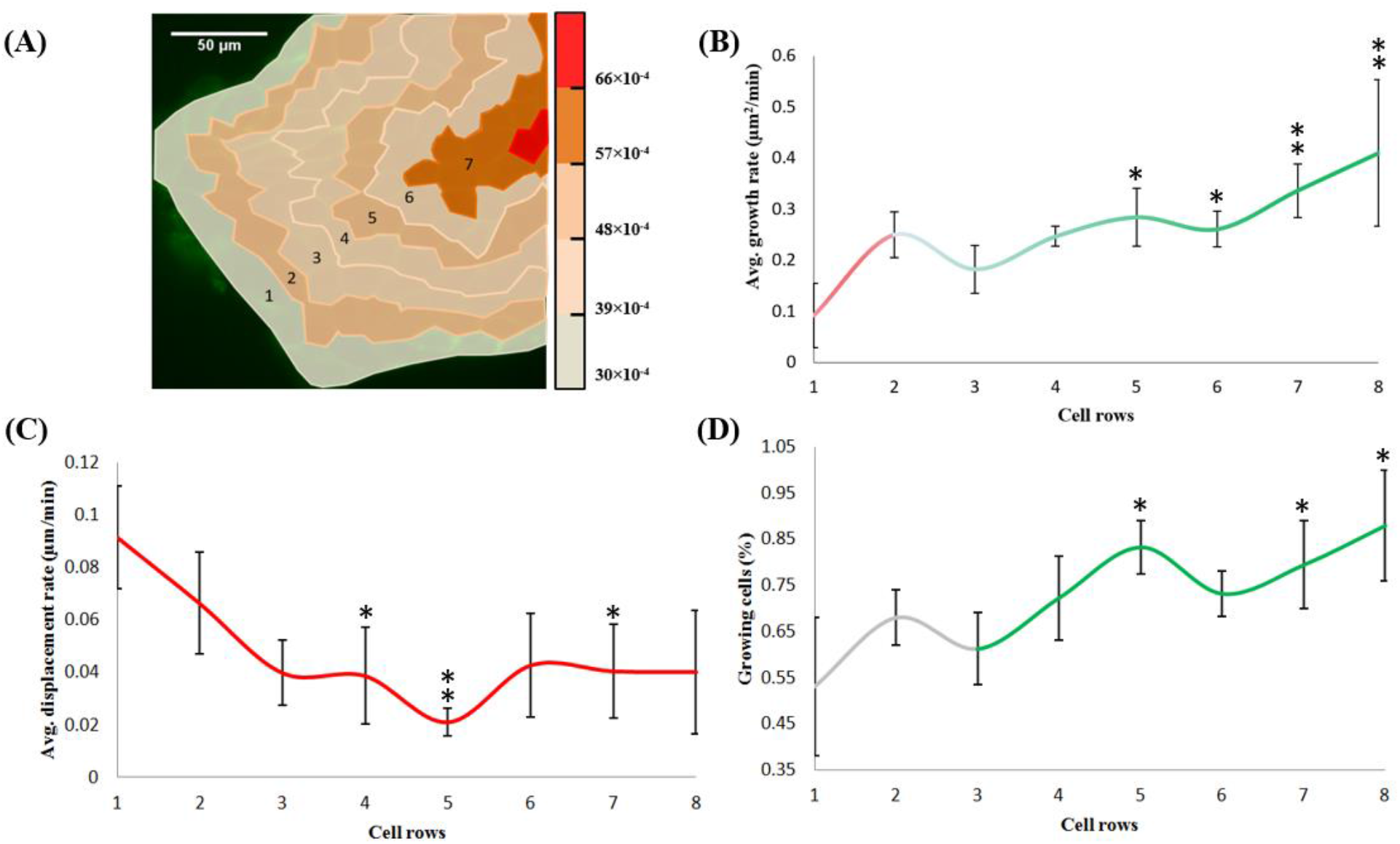
Growing cell colonies backup the migration of the leader cells (A) Average strain, (B) average growth rate, (C) average displacement rate, (D) percentage of growing cell distribution among various cell rows, (E) collective cell migration model. Note: Red color indicates displacement, and green color indicates growth dominant regions. Note: Error bars in (B), (C), and (D) represents the standard error obtained from 5 independent experiments. These analyses were done on static cell colonies.

It was found that GR increased along the cell rows (Figure 8B) because expanding (or growing) cells existed in the sub-marginal region. Whereas the DR decreased (Figure 8C) along the cell rows as the leader cells had unconstrained boundaries, they can migrate at a higher rate than the follower cells. It was observed that GR (Figure 8B) and DR (Figure 8C) showed the opposite trend. Thereby, comparing the GR and DR along the rows exhibited a negative Pearson correlation (r = −0.64). It implies that rows with a high growth rate undergo less displacement and vice versa. However, the GR (Figure 8B) along the cell rows exhibited a similar trend as that of the percentage of growing cells (Figure 8D). The sudden rise in the GR observed in the 2^nd^ row (Figure 8B) was due to their stretching caused due to the pull generated by the leader cells (Figure 7). Thereby, a high strain (S) was seen in the 2^nd^ row (Figure 8A). Further, high strain (Figure 8A), peak in GR (Figure 8B), and dip in DR (Figure 8C) were observed in the 5^th^ cell row. Interestingly, the 5^th^ row also contained a high percentage of growing cells (Figure 8D). It suggests that growing cells displace less. Similarly, moving along the cell rows, the S and GR varied depending upon the number of growing cells available in that row. The statistical analysis was carried out using repeated ANOVA to determine the significant difference of GR, DR, and percentage of growing cells among the rows. The GR (Figure 8B) of 1^st^ row was significantly lesser than 5^th^ row (p = 0.023), 6^th^ row (p = 0.044), 7^th^ row (p = 0.005) and 8^th^ row (p = 0.002). The DR (Figure 8C) of the 1^st^ row was significantly higher than the 4^th^ row (p = 0.043), 5^th^ row (p = 0.009) and 7^th^ row (p = 0.045). The percentage of growing cells (Figure 8D) in the 5^th^ row (p = 0.024), 7^th^ row (p = 0.044) and 8^th^ row (p = 0.002) was significantly higher as compared to the 1^st^ row.

### 3.5 Growing colonies during longer experiment

Till now, we have observed growing cells colonies in up to 4 hours. To check their existence after 4 hours, we conducted 12 hr experiment. The 12 hr experiment showed the presence of huge, medium, and small colonies for 0-4 hr (Figure 9A), 4-8 hr (Figure 9B), and 8-12 hr (Figure 9C), suggesting the predominance of the growing colony at an early stage of migration (0-4 hr). Interestingly, the percentage expansion in the average cell area (Figure 9D) and average velocity (Figure 9E) varied in accordance with the size (number of cells) of the colonies (Figure 9A-C). The small size colonies were observed from 8-12 hr (Figure 9C). The possible reason for this could be the high average cell area, i.e., 243 μm^2^ at t = 8 hr. Thereby, the majority of cells were unable to quality the 20% growth criteria. We assume that after this point, the cells of the leading site become purely motile cells without much fluctuation in their area.

**Figure 9.**
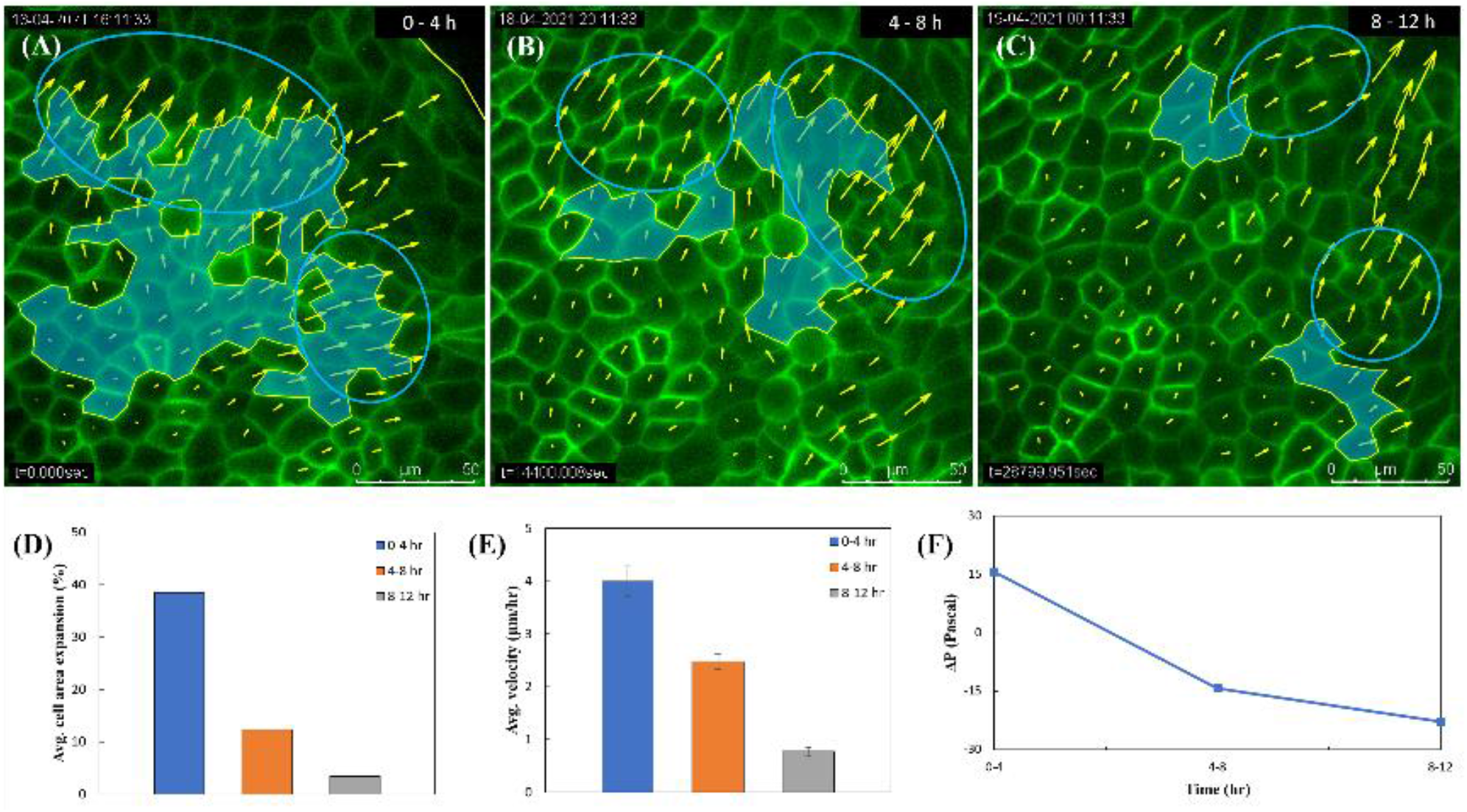
The 12-hour experiment at the leading site (A) Growing colony from 0-4 hr, (B) growing colonies from 4-8 hour, (C) growing colonies from 8-12 hr, (D) average cell area expansion (%), (E) Average velocity, and (F) the ΔP values from 0-4, 4-8 and 8-12 hr. Note: These analyses were done on static cell colonies. The blue ellipse shows the highly aligned velocity field.

Further, we computed the Δ***P*** of the monolayer. Our results showed that at an early stage of migration (0-4 hr), the monolayer was under pressure (+Δ***P***) in the presence of a big colony (Figure 9F). However, later stage of migration, from 4-8 hr, the transition occurred in the monolayer from being under pressure (+Δ***P***) to tension (-Δ***P***) in the presence of medium colonies (Figure 9F). At last, from 8-12 hr, the tension of the monolayer increased in the presence of smaller colonies (Figure 9F). It implies that in an early stage of migration (0-4 hr), the leading site of the monolayer is under pressure due to the presence of a big colony that is capable of generating enough pressure (compressive force) to promote the formation of the leader cells. Later, the colony expands, thereby displacing the front rows towards the leader cells to exhibit collective cell migration (Figure S3A, Figure 4A, Figure S9A and Figure 9A). However, these colonies start to disappear in the later stages of migration (4-8 hr (Figure 9B) and 8-12 hr (Figure 9C)), increasing the monolayer tension. Therefore, our study indicates that the compressive force applied by the colonies might be responsible for leader cell recruitment.

Finally, as per our understanding, we propose a model for the expansion of the epithelial cell monolayer (Figure S5E). Initially, from 0-4 hr, the cell expansion dominates the migration, as huge expanding colonies are observed in the sub-marginal region. These colonies apply localized compressive force on their neighbouring cells that may be sensed by some of the marginal cells, and they get transformed into leader cells. Then, these newly recruited leader cells pull their immediate follower cells or front cell rows (Figure S5D and Figure 7) that are already being pushed by colonies. Thereby, resulting in the displacement of front cell rows towards the leader cells. Therefore, highly aligned collective cell motion is observed at the leading cell rows (shown by a blue ellipse in Figure S3A and Figure 4A). The cells of the colonies spread to cover the gap produced by the displacement of front cell rows in order to maintain tissue integrity. Thereby, the colonies expand or spread towards the leader cells (Figure S3C and Figure 4B). However, later, from 4-12 hr, the expanding colonies start to disappear, suggesting the dominance of cell migration over expansion.

## 4 Discussion

In this work, we aimed to identify the role of follower cells in collective cell migration of freely expanding monolayer. For simplification, it was assumed that the intercellular forces acting within the cell-cell junction dominates the epithelial mechanics (Vincent et al., 2013). Here, we observed that a few groups of cells were expanding at a higher rate than the rest of the tissue. To understand the effect of this differential growth rate in the monolayer, we referred to the theoretical model given by Boris I. Shraiman (Shraiman, 2005). He explained the presence of a feedback mechanism that could be responsible for regulating the tissue growth in the wings of the Drosophila melanogaster. His model focused on the centre of a tissue where cell number density is quite high. He assumed that cells behave like solid and possess high rigidity to shear. He further analyzed the concept of “cell competition,” where a group of cells growing with a higher rate, called winner cells, push their slow-growing neighbouring cells, called loser cells. This inhomogeneous growth rate leads to the generation of localized mechanical stress in the monolayer, which gets relieved by the elimination of the loser cells via apoptosis (Shraiman, 2005; Vincent et al., 2013; Matamoro-Vidal and Levayer, 2019). The winner cells being highly sensitive, immediately sense the apoptotic factor released by the dying cell (Tsuboi et al., 2018). Thereby, the winner cells expand to fill the space generated by the collapse of the dying cells to maintain tissue integrity. Hence this promotes the biased area growth of the winner cells (Tsuboi et al., 2018). Boris I. Shraiman (Shraiman, 2005) claimed that his model is of general relevance and can be applied where the differential growth rate exists among the cell population.

In our experimental study, we found that there was no colony at the centre (far away from the leading edge) of the monolayer (Figure S8A). However, the colonies start to appear after the events of apoptosis. Later, the number of cells forming these colonies increased with the events of apoptosis (Figure S8B-D). The apoptosis may have occurred to relieve mechanical stress. Thereby, few winner cells located in the vicinity of the dying cell grow in size to fill the space vacated by it. These winner cells together represented a colony. Hence, the colonies emerged due to the events of apoptosis. Whereas, at the leading site, we assumed that the cells behave like viscoelastic material with certain rigidity to shear, and a finite number of cell-cell rearrangements can occur. As indicated by Boris I. Shraiman’s theoretical model (Shraiman, 2005), we experimentally observed two types of cell populations at the leading site, classified as winners and losers. The winners gain in area, and losers lose in their area due to the cell competition. In our study, the colonies’ cells were considered winner cells, and surrounding cells were considered loser cells. The average growth rate of winner cells was significantly (p = 0.0007) higher as compared to the loser cells (Figure 10A). Interestingly, the initial average area of winner cells was less than the loser cells (Figure 10B). However, then winners grew at a significantly higher rate than the losers. Therefore, with time, the average area of winners exceeds the losers (Figure 10B). Interestingly, at every time step, the value of MSD for losers was more than winners (Figure 10C), suggesting that loser cells displace more than the winner cells. It means that winner cells displace less and loser cells displace more.

**Figure 10.**
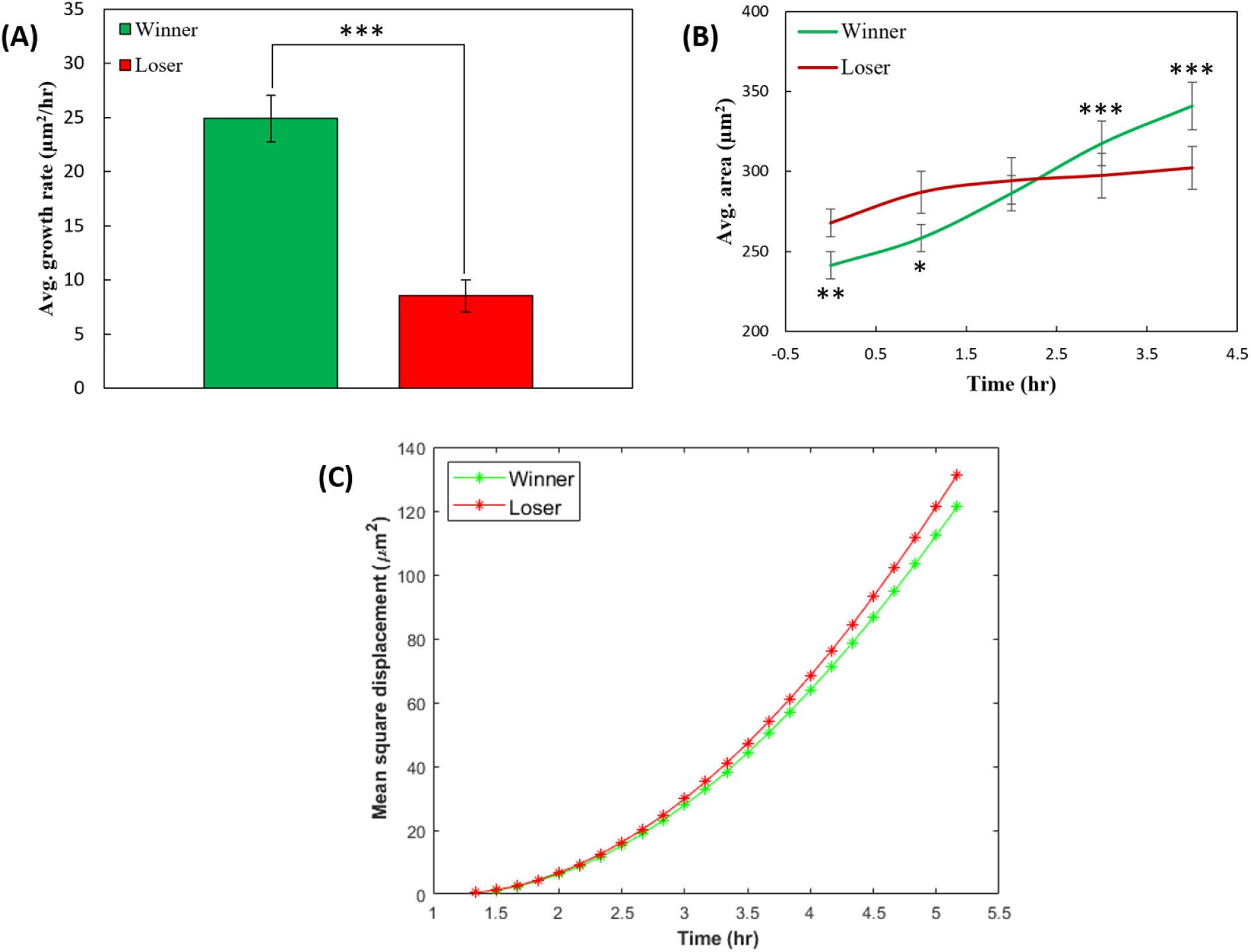
Classification of winner and loser cells (A) Average areal growth rate, (B) average area, and (C) mean square displacement as a function of time. Note: The error bar represents standard error from 5 independent experiments. The winner cells were compared with loser cells. These analyses were done on static cell colonies.

Some mutations induce a sudden high growth rate in the cells of the expanding colonies (Shraiman, 2005). These mutations only affect the growth rate of the cells in real-time, without changing the division rate (follow regular cell cycle time). The growth rate of the cells of colonies is much higher as compared to the regular division rate observed in the monolayer. Hence, cell division has a minor effect on the mechanism discussed here. During our experiments, very few cell divisions were observed, which were excluded in this study. Another reason for neglecting the dividing cell was that the collective cell migration does not stop in the absence of cell division (Gauquelin et al., 2019). The differential growth rate among the growing (or winner) cells and remaining (or loser) cells are responsible for cell competition. With time, the area of growing or winner cells becomes larger than the neighbouring or loser cells (Figure 10B). Stefan et al. (Nehls et al., 2019) reported that the stiffness of MDCK cells depends upon their size, i.e., big cells have high stiffness, and small cells have less stiffness. Therefore, it suggests that the winner cells of the colonies have higher stiffness compared to the loser cells. As per the literature, the differential growth rate and high motility found in the MDCK monolayer make it an optimum system where cell competition can occur (Basan et al., 2009, 2011; Kon et al., 2017). Here, we experimentally showed the effect of cell competition at the leading site of the MDCK cell monolayer. Where colonies present at the leading site have high homeostatic pressure (or low tension), push their front tissue having low pressure (or high tension). We assumed that some of the marginal cells sense this push and transform into the leader cells. These leader cells start pulling their immediate neighbours present in the 2^nd^ row. Meanwhile, the cells of the colonies expand in size, thereby displacing the front rows towards the leader cells. Therefore, the net force generated due to the cell competition among the follower cells acts in a similar direction in which the leaders were pulling the monolayer. Thereby, both the force might be involved collectively in displacing the front rows towards the leader cells.

In the 12 hour experiments (Figure 9), the colonies start to disappear in the later stage of migration (4-12 hr), the formation of colonies can be linked to backward propagating slow mechanical waves (Serra-Picamal et al., 2012; Tlili et al., 2018; Boocock et al., 2021). However, the focus of this work was to investigate the roles of the growing colonies in the recruitment of leader cells and collective cell migration observed in the front rows. Boris et al. showed that a patch or group of winner cells growing at a higher rate than surrounding tissue could cause the generation of localized mechanical stress that is compressive (Shraiman, 2005). Janet et al. have demonstrated that compressive stress can enhance the migration and promote the formation of the leader cells at the wound periphery (Janet et al., 2012). Similarly, in our case, we assume that expanding colonies generates localized compressive stress in the submarginal region that could be responsible for recruiting the leader cells. Reffay et al. demonstrated that upon inhibition of RhoA in migrating epithelial monolayer, the leader cells were being pushed by the follower cells (Reffay et al., 2014). It suggests the presence of an alternative mechanism wherein the absence of the leader cells, the pushing forces generated by the sub-marginal cells promotes the migration of complete monolayer. Since the growing colonies could be the source of the push originated from the follower cells, it suggests that the local expansion of colonies can back up the migration of front rows in the absence of leader cells. In our case, we observed that the velocity vectors were highly aligned near the head of colonies, pointing towards the front cell rows, even when the size of colonies was small (Figure S9 and Figure 9A-C). Therefore, it was assumed that the push from the growing colonies and pull from the leader cells collectively displace the front cell rows. It could be the reason for the highly aligned velocity field observed at the head of the colonies and in the front rows (Figure S3A, Figure 4A, Figure S9, and Figure 9A-C). The transition of monolayer from being under compression (+Δ***P***) at the early stage (0-4 hr) to under tension (−Δ***P***) at the later stage of migration (8-12 hr) was observed. It could be the reason for the predominantly tensile stress reported by Xavier et al. for migrating epithelial monolayer (Trepat et al., 2009a).

In literature, there exist several physical and chemical mechanisms to explain collective cell migration. The coordination of CIL and local alignment (LA) has been reported to result in collective cell migration. The coordination of CIL and LA has been shown to dictate giant density fluctuations in the monolayer, resulting in phase separation (Smeets et al., 2016; Lin et al., 2018, 2019). In our case, winner cells (of the colonies) grow at a higher rate than the loser cells (of the surrounding tissue), which could be treated as phase separation. It is reported in the literature that when LA dominates CIL, then translation motion occurs, and when CIL dominates LA, then cage-relative motion occurs. The high MSD of loser cells compared to winner cells (Figure 10C) suggest that LA dominates in surrounding tissue and CIL dominates inside the colonies (Kiran et al., 2021). Further, Jain et al. (Jain et al., 2020) discussed that the winner cell has less slope than the loser cell. Thereby, the lamellipodia of the winner cell undergo beneath the loser cell and re-polarizes it (loser cell) towards its (winner cell) direction of motion (Jain et al., 2020). In our case, the winner cells of the colonies spread (increase in area and decrease in height) with time resulting in less height compared to surrounding loser cells (Figure 6D-E). Therefore, it can be assumed that winner cells of the colonies might re-polarize their neighbouring loser cells in the direction of their spreading, resulting in the highly aligned velocity field at the head of the colony and front rows (Figure S3A, Figure 4A, Figure S9A and Figure 9A). Some chemical mechanism (e.g., activation of Rho GTPases, intracellular dynamics of Merlin) might also be involved in this case which is a scope of future investigation.

## 5 Conclusions

The expansion of the epithelial cell monolayer is an essential biological phenomenon that occurs in nature. The collective cell migration is an important feature observed in the expansion of the epithelial tissues. Many studies have suggested that the follower cells also play a critical role in collective cell migration (Farooqui and Fenteany, 2005; Trepat et al., 2009b; Petitjean et al., 2010; Tambe et al., 2011; Serra-Picamal et al., 2012). However, the physical mechanism through which follower cells contribute to the expansion of the epithelial tissue is not well known. Therefore, our study aimed to identify the mechanism through which follower cells coordinate with leader cells to achieve collective cell motion in freely expanding epithelial tissue. We have found the presence of dynamically expanding cell colonies in the sub-marginal region. Although there was no significant difference (p = 0.6347) between the population (%) of the colonies cells (n = 53%) and the remaining cells (n = 47%), the contribution of the colonies cells (a = 81%) in the overall expansion of the leading site was statistically significant (p = 0.0007) as compared to the remaining cells (a = 19%). Moreover, the expansion ratio of the colonies was statistically significant (p < 0.05) to the monolayer (colonies and remaining cells combined). It confirms that these expanding colonies majorly contribute to the net expansion of the leading site. We assumed that these expanding colonies might push their neighbouring cells. The 3D analysis was done to find the real meaning of expanding colonies and trace their push to the neighbouring cells. We found that the cells of the colonies were increasing in their area and decreasing in height. In comparison, the surrounding cells shrink in their area and have more height than the cells of the colonies. Then we superimposed the initial and final shape of the static colonies. It was found that these colonies expand towards the leader cells. Thereby, we hypothesized that these expanding colonies apply maximum push towards the marginal cells. It may trigger some of the marginal cells, and they transform into the leader cells. These newly recruited leader cells start pulling their follower cells (or front rows) that are already being pushed by spreading colonies, resulting in the displacement of front rows towards the leader cell. It may be the reason for the collective cell migration observed in the front rows. Perhaps, this could be the plausible mechanism through which follower cells contribute to the collective cell migration in freely expanding epithelial monolayer. However, further investigation is required for a better understanding of this mechanism.

## 7 Author Contributions

Abhimanyu Kiran designed and performed the experiments, did the analysis and interpretation of data, written the original manuscript. Navin Kumar and Vishwajeet Mehandia edited and drafted the final version of the manuscript.

## 9 Data Accessibility

The datasets supporting this article have been uploaded as part of the Supplementary Material. We thank W. J. Nelson for sending the MDCK cell lines.

## 10 Acknowledgement

We thank W. J. Nelson for sending the MDCK cell lines.

## References

Basan, M., Prost, J., Joanny, J.-F., and Elgeti, J. (2011). Dissipative particle dynamics simulations for biological tissues: rheology and competition. Phys. Biol. 8, 26014.

Basan, M., Risler, T., Joanny, J.-F., Sastre-Garau, X., and Prost, J. (2009). Homeostatic competition drives tumor growth and metastasis nucleation. HFSP J. 3, 265–272.

Boocock, D., Hino, N., Ruzickova, N., Hirashima, T., and Hannezo, E. (2021). Theory of mechanochemical patterning and optimal migration in cell monolayers. Nat. Phys. 17, 267–274.

Coburn, L., Lopez, H., Caldwell, B. J., Moussa, E., Yap, C., Priya, R., et al. (2016). Contact inhibition of locomotion and mechanical cross-talk between cell--cell and cell--substrate adhesion determine the pattern of junctional tension in epithelial cell aggregates. Mol. Biol. Cell 27, 3436–3448.

Du Roure, O., Saez, A., Buguin, A., Austin, R. H., Chavrier, P., Siberzan, P., et al. (2005). Force mapping in epithelial cell migration. Proc. Natl. Acad. Sci. 102, 2390–2395.

Farooqui, R., and Fenteany, G. (2005). Multiple rows of cells behind an epithelial wound edge extend cryptic lamellipodia to collectively drive cell-sheet movement. J Cell Sci 118, 51–63.

Friedl, P., and Gilmour, D. (2009). Collective cell migration in morphogenesis, regeneration and cancer. Nat. Rev. Mol. Cell Biol. 10, 445–457. doi:10.1038/nrm2720.

Friedl, P., Hegerfeldt, Y., and Tusch, M. (2004). Collective cell migration in morphogenesis and cancer. Int. J. Dev. Biol. 48, 441–449.

Gauquelin, E., Tlili, S., Gay, C., Peyret, G., Mège, R.-M., Fardin, M. A., et al. (2019). Influence of proliferation on the motions of epithelial monolayers invading adherent strips. Soft Matter 15, 2798–2810.

Gov, N. S. (2007). Collective cell migration patterns: follow the leader. Proc. Natl. Acad. Sci. 104, 15970–15971.

Ilina, O., and Friedl, P. (2009). Mechanisms of collective cell migration at a glance. J. Cell Sci. 122, 3203–3208. doi:10.1242/jcs.036525.

Jain, S., Cachoux, V. M. L., Narayana, G. H. N. S., de Beco, S., D’alessandro, J., Cellerin, V., et al. (2020). The role of single-cell mechanical behaviour and polarity in driving collective cell migration. Nat. Phys. 16, 802–809.

Janet, M. T., Cheng, G., Tyrrell, J. A., Wilcox-Adelman, S. A., Boucher, Y., Jain, R. K., et al. (2012). Mechanical compression drives cancer cells toward invasive phenotype. Proc. Natl. Acad. Sci. 109, 911–916.

Jasaitis, A., Estevez, M., Heysch, J., Ladoux, B., and Dufour, S. (2012). E-cadherin-dependent stimulation of traction force at focal adhesions via the Src and PI3K signaling pathways. Biophys. J. 103, 175–184.

Kiran, A., Kumar, N., and Mehandia, V. (2021). Distinct Modes of Tissue Expansion in Free Versus Earlier-Confined Boundaries for More Physiological Modeling of Wound Healing, Cancer Metastasis, and Tissue Formation. ACS Omega.

Kon, S., Ishibashi, K., Katoh, H., Kitamoto, S., Shirai, T., Tanaka, S., et al. (2017). Cell competition with normal epithelial cells promotes apical extrusion of transformed cells through metabolic changes. Nat. Cell Biol. 19, 530–541.

Ladoux, B. (2009). Biophysics: cells guided on their journey. Nat. Phys. 5, 377.

Lin, S.-Z., Bi, D., Li, B., and Feng, X.-Q. (2019). Dynamic instability and migration modes of collective cells in channels. J. R. Soc. Interface 16, 20190258.

Lin, S.-Z., Ye, S., Xu, G.-K., Li, B., and Feng, X.-Q. (2018). Dynamic migration modes of collective cells. Biophys. J. 115, 1826–1835.

Magno, R., Grieneisen, V. A., and Marée, A. F. M. (2015). The biophysical nature of cells: potential cell behaviours revealed by analytical and computational studies of cell surface mechanics. BMC Biophys. 8, 1–37.

Maruthamuthu, V., Sabass, B., Schwarz, U. S., and Gardel, M. L. (2011). Cell-ECM traction force modulates endogenous tension at cell--cell contacts. Proc. Natl. Acad. Sci. 108, 4708–4713.

Matamoro-Vidal, A., and Levayer, R. (2019). Multiple Influences of Mechanical Forces on Cell Competition. Curr. Biol. 29, R762--R774.

Mayor, R., and Etienne-Manneville, S. (2016). The front and rear of collective cell migration. Nat. Rev. Mol. cell Biol. 17, 97.

Mertz, A. F., Banerjee, S., Che, Y., German, G. K., Xu, Y., Hyland, C., et al. (2012). Scaling of traction forces with the size of cohesive cell colonies. Phys. Rev. Lett. 108, 198101.

Mertz, A. F., Che, Y., Banerjee, S., Goldstein, J. M., Rosowski, K. A., Revilla, S. F., et al. (2013). Cadherin-based intercellular adhesions organize epithelial cell--matrix traction forces. Proc. Natl. Acad. Sci. 110, 842–847.

Nehls, S., Nöding, H., Karsch, S., Ries, F., and Janshoff, A. (2019). Stiffness of MDCK II Cells Depends on Confluency and Cell Size. Biophys. J. 116, 2204–2211.

Notbohm, J., Banerjee, S., Utuje, K. J. C., Gweon, B., Jang, H., Park, Y., et al. (2016). Cellular contraction and polarization drive collective cellular motion. Biophys. J. 110, 2729–2738.

Petitjean, L., Reffay, M., Grasland-Mongrain, E., Poujade, M., Ladoux, B., Buguin, A., et al. (2010). Velocity fields in a collectively migrating epithelium. Biophys. J. 98, 1790–1800.

Poujade, M., Grasland-Mongrain, E., Hertzog, A., Jouanneau, J., Chavrier, P., Ladoux, B., et al. (2007). Collective migration of an epithelial monolayer in response to a model wound. Proc. Natl. Acad. Sci. 104, 15988–15993.

Reffay, M., Parrini, M.-C., Cochet-Escartin, O., Ladoux, B., Buguin, A., Coscoy, S., et al. (2014). Interplay of RhoA and mechanical forces in collective cell migration driven by leader cells. Nat. Cell Biol. 16, 217.

Rørth, P. (2009). Collective Cell Migration. Annu. Rev. Cell Dev. Biol. 25, 407–429. doi:10.1146/annurev.cellbio.042308.113231.

Serra-Picamal, X., Conte, V., Vincent, R., Anon, E., Tambe, D. T., Bazellieres, E., et al. (2012). Mechanical waves during tissue expansion. Nat. Phys. 8, 628.

Shraiman, B. I. (2005). Mechanical feedback as a possible regulator of tissue growth. Proc. Natl. Acad. Sci. 102, 3318–3323.

Smeets, B., Alert, R., Pešek, J., Pagonabarraga, I., Ramon, H., and Vincent, R. (2016). Emergent structures and dynamics of cell colonies by contact inhibition of locomotion. Proc. Natl. Acad. Sci. 113, 14621–14626.

Szabo, B., Szöllösi, G. J., Gönci, B., Jurányi, Z., Selmeczi, D., and Vicsek, T. (2006). Phase transition in the collective migration of tissue cells: experiment and model. Phys. Rev. E 74, 61908.

Tambe, D. T., Hardin, C. C., Angelini, T. E., Rajendran, K., Park, C. Y., Serra-Picamal, X., et al. (2011). Collective cell guidance by cooperative intercellular forces. Nat. Mater. 10, 469.

Tlili, S., Gauquelin, E., Li, B., Cardoso, O., Ladoux, B., Delanoë-Ayari, H., et al. (2018). Collective cell migration without proliferation: density determines cell velocity and wave velocity. R. Soc. open Sci. 5, 172421.

Trepat, X., Wasserman, M. R., Angelini, T. E., Millet, E., Weitz, D. A., Butler, J. P., et al. (2009a). Physical forces during collective cell migration. Nat. Phys. 5, 426–430. doi:10.1038/nphys1269.

Trepat, X., Wasserman, M. R., Angelini, T. E., Millet, E., Weitz, D. A., Butler, J. P., et al. (2009b). Physical forces during collective cell migration. Nat. Phys. 5, 426.

Tsuboi, A., Ohsawa, S., Umetsu, D., Sando, Y., Kuranaga, E., Igaki, T., et al. (2018). Competition for space is controlled by apoptosis-induced change of local epithelial topology. Curr. Biol. 28, 2115–2128.

Vedula, S. R. K., Leong, M. C., Lai, T. L., Hersen, P., Kabla, A. J., Lim, C. T., et al. (2012). Emerging modes of collective cell migration induced by geometrical constraints. Proc. Natl. Acad. Sci. 109, 12974–12979.

Vicsek, T., Czirók, A., Ben-Jacob, E., Cohen, I., and Shochet, O. (1995). Novel type of phase transition in a system of self-driven particles. Phys. Rev. Lett. 75, 1226.

Vincent, J.-P., Fletcher, A. G., and Baena-Lopez, L. Al. (2013). Mechanisms and mechanics of cell competition in epithelia. Nat. Rev. Mol. cell Biol. 14, 581–591.

Vishwakarma, M., Di Russo, J., Probst, D., Schwarz, U. S., Das, T., and Spatz, J. P. (2018). Mechanical interactions among followers determine the emergence of leaders in migrating epithelial cell collectives. Nat. Commun. 9, 1–12.

Vitorino, P., and Meyer, T. (2008). Modular control of endothelial sheet migration. Genes Dev. 22, 3268–3281.

Zehnder, S. M., Suaris, M., Bellaire, M. M., and Angelini, T. E. (2015). Cell volume fluctuations in MDCK monolayers. Biophys. J. 108, 247–250.

Zimmermann, J., Camley, B. A., Rappel, W.-J., and Levine, H. (2016). Contact inhibition of locomotion determines cell--cell and cell--substrate forces in tissues. Proc. Natl. Acad. Sci. 113, 2660–2665.

