## Supplementary figures for "Mechanism of tissue expansion in the early stage of the migration"

### *Supplementary Material*

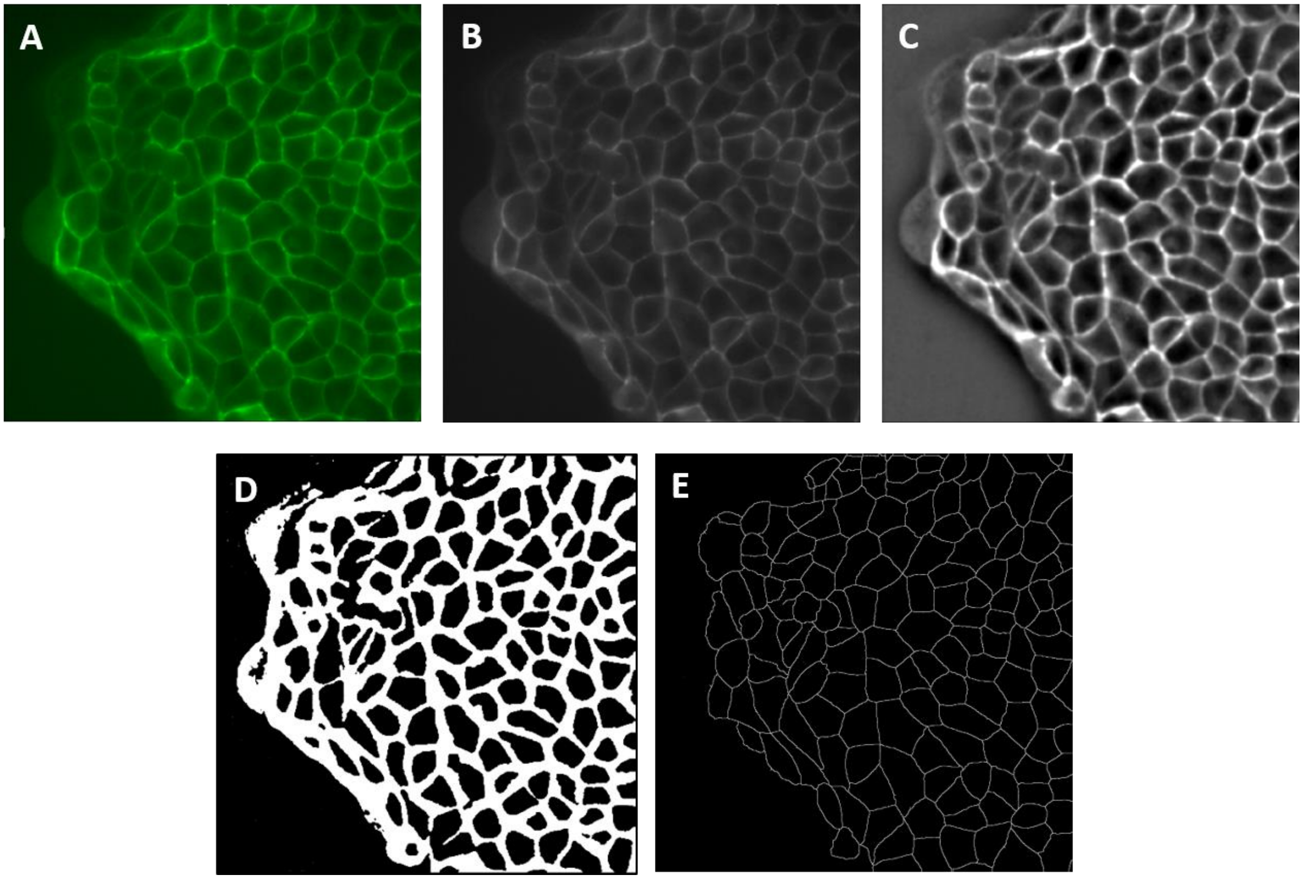


**Figure S1.** Segmentation of image sequence (A) Load image sequence in ImageJ, (B) convert it into 8-bit images, (C) apply bandpass FFT filter, (D) convert images into binary format, and (E) skeletonization of the images.

<u>Protocol for the segmentation of image sequence</u>

1. Open ImageJ and load raw image sequence in the ImageJ (Fig. S1a)

    File > Import > Image Sequence

2. Convert the raw images to the 8-bit images (Fig. S1b).

    Image > Type > 8-bit

3. Apply bandpass Fast Fourier transform (FFT) filter (Fig. S1c)

    Process > FFT> Bandpass Filter
    ok

4. Convert it into binary images (Fig. S1d).

Process > Binary > Make Binary
Select method 'Percentile' and background 'Light'
Tick checkbox 'Calculate threshold for each image'
ok

5. Segmentation of the images (Fig. S1e).

Process > Binary > Skeletonize

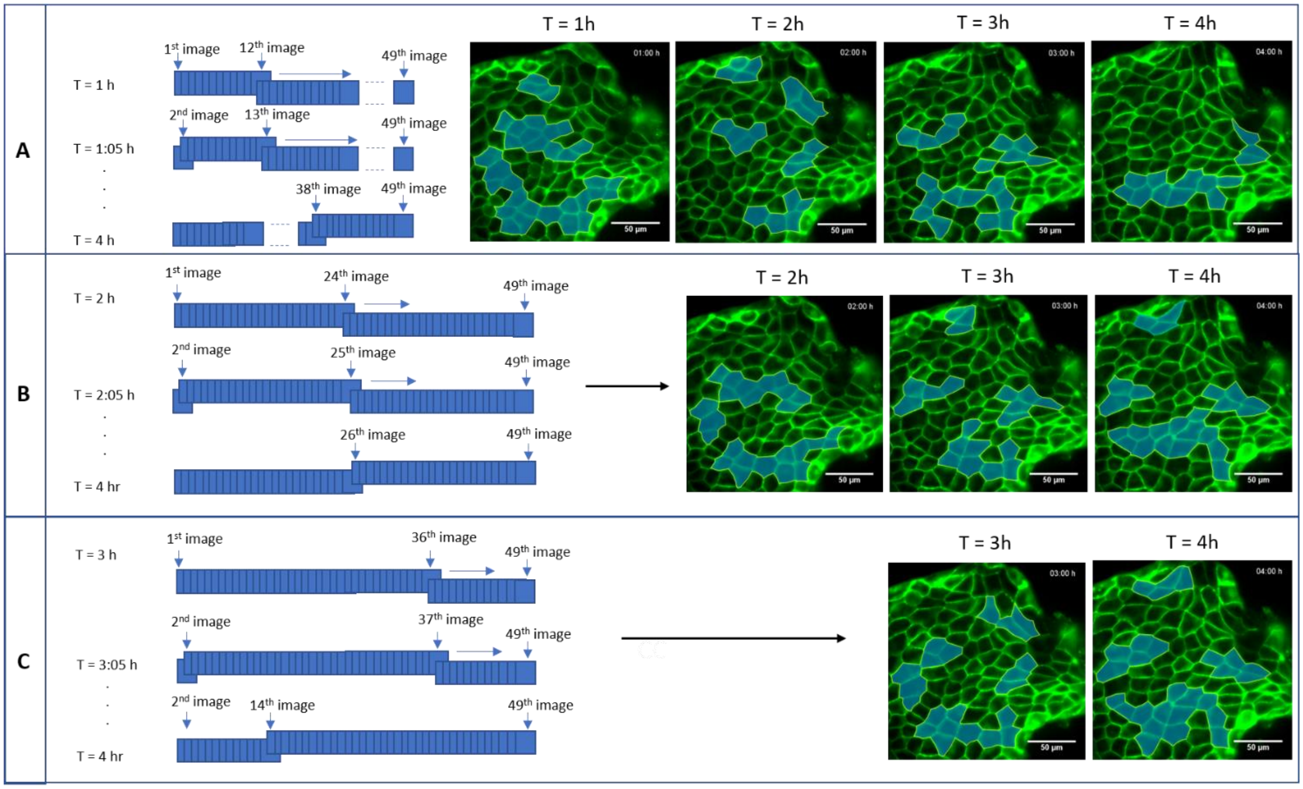


**Figure S2.** Moving window analysis by varying the window size (A) 12 images (or 1 hr), (B) 24 images (or 2 hr), and (C) 36 images (or 3 hr).

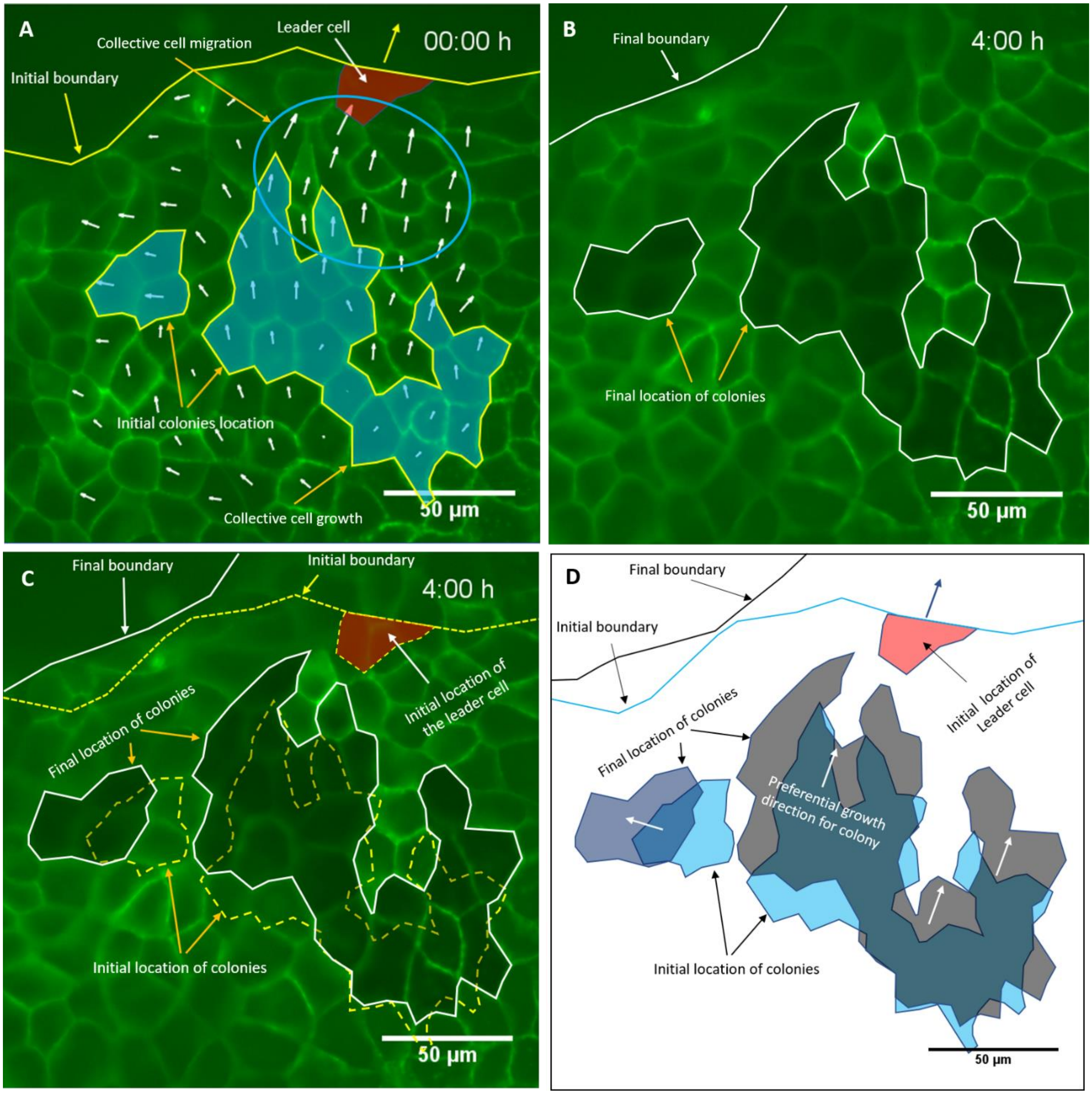


**Figure S3.** The colonies found near the margin prefer to grow towards the leader cells (A) Initial shape and location of the colonies at t = 0 h with velocity vectors obtained from initial and final cell positions, (B) final shape and location of the colonies, (C) super positioning of the initial location and final location of the colonies, (D) analysis of the preferential growth direction for the colonies. Note: The static growing colonies were considered for this analysis.

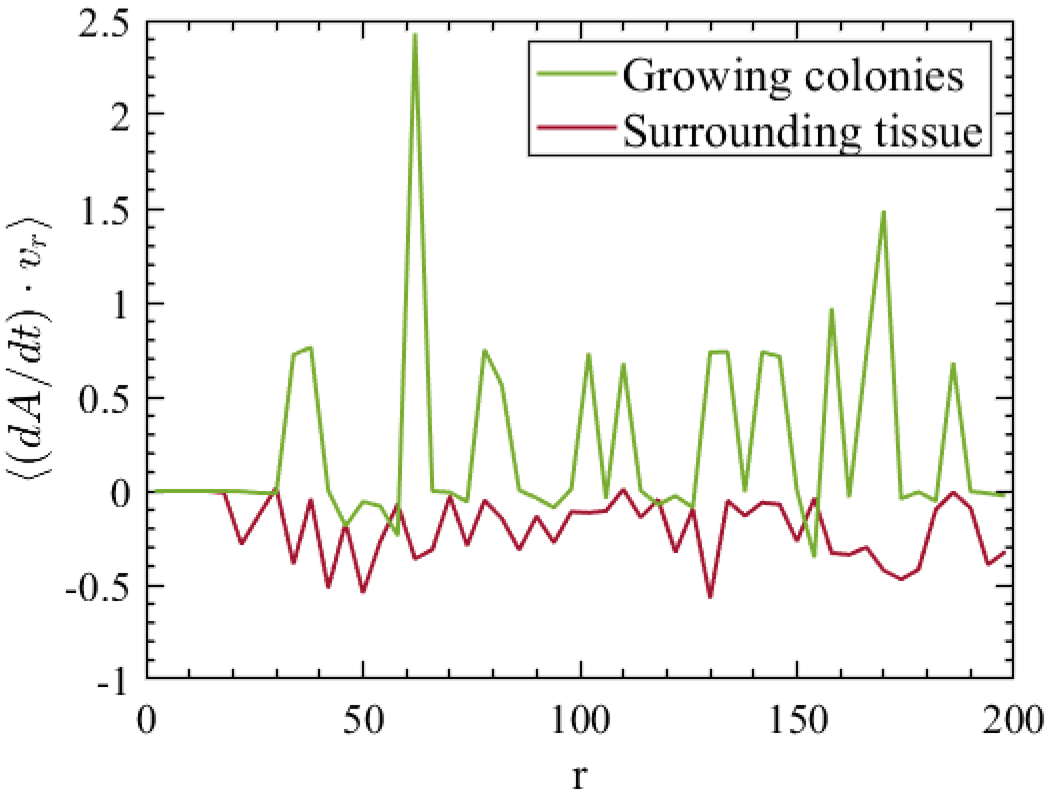


**Figure S4.** The plot of growth rate-velocity correlation as a function of space for growing colonies and surrounding tissue.

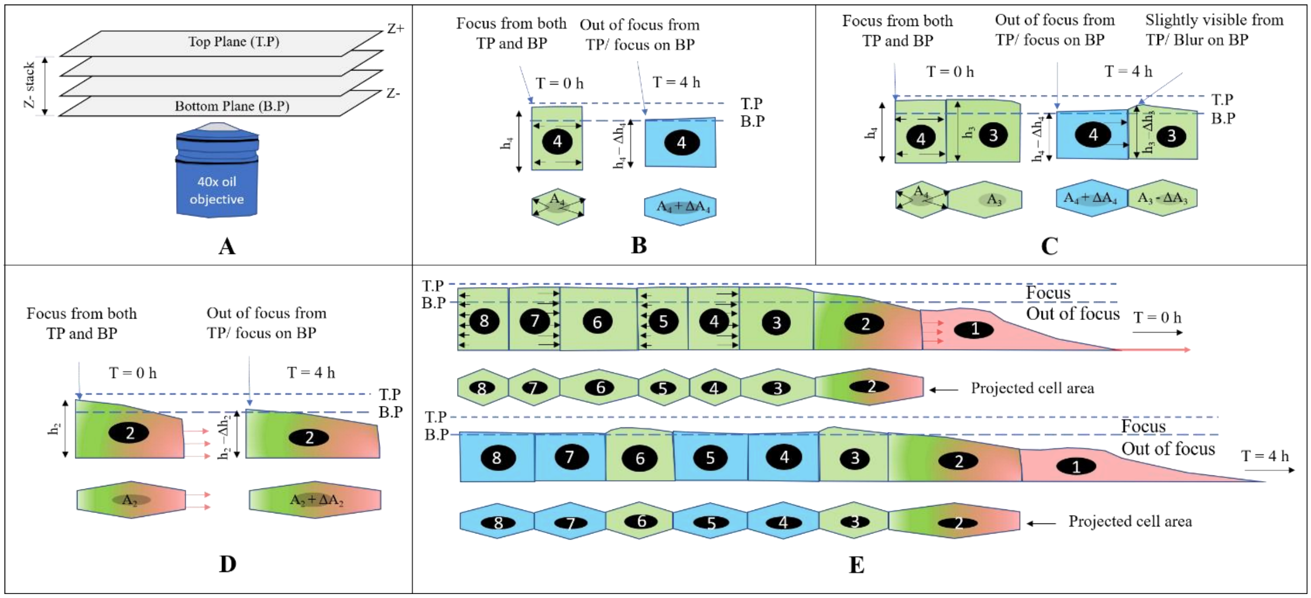


**Figure S5.** The analysis of z-stack images (A) Schematic of Z-stack imaging, (B) cell stretching, (C) expanding cell pushing their neighboring cells, (D) cell stretching, and (E) collective cell migration. Note: Blue color represents expanding (or spreading) colony cell.

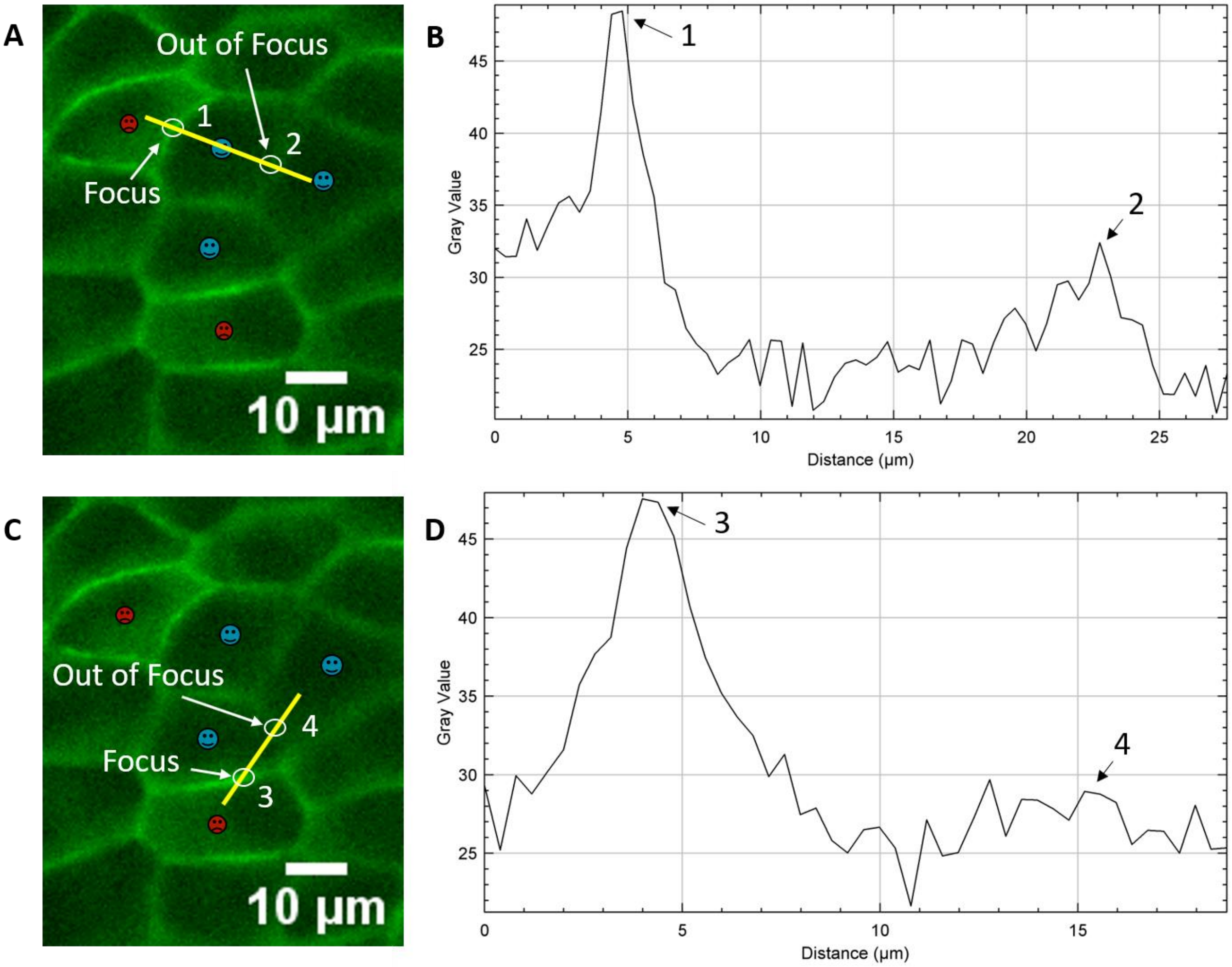


**Figure S6.** Method of identification of focus and out of focus regions (A) The image showing focus edge 1 and out of focus edge 2, (B) The gray-scale values across the line profile passing through edge 1 and 2, (C) Image containing focus edge 3 and out of focus edge 4, (D) The gray-scale values across the line profile passing through edge 3 and 4. Note: The Red and blue faces represent shrinking and expanding cells, respectively. The static growing colonies were considered for this analysis.

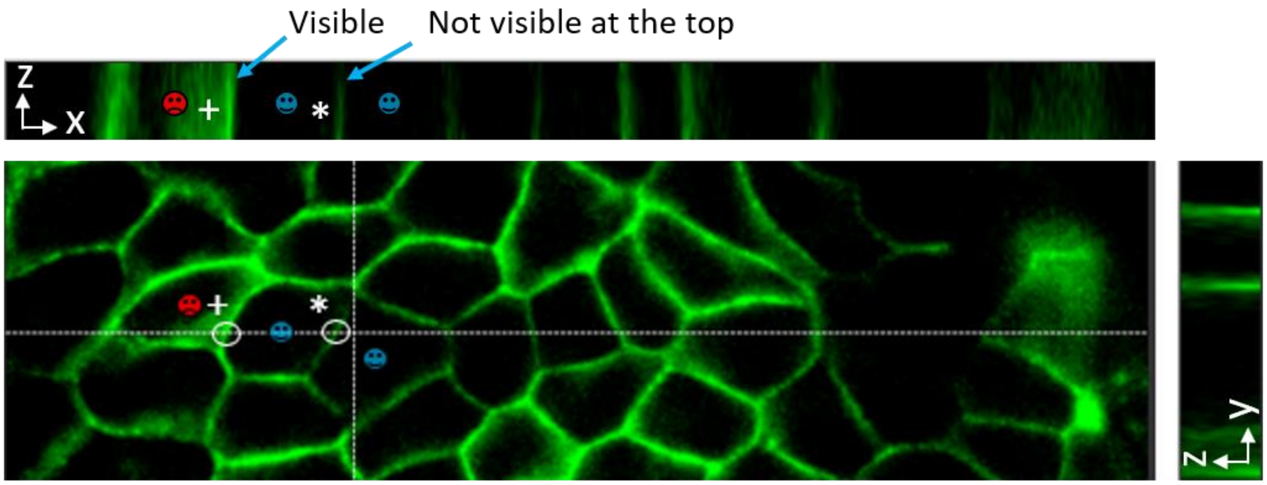


**Figure S7.** The orthogonal view of the shrinking and expanding cells. The common edge of the shrinking and expanding cell was visible (represented by symbol +) throughout the z-stack. The common edge of expanding cells was not visible (represented by symbol *) on the top of the z-stack. Note: Red and blue faces represent shrinking and expanding cells, respectively. The static growing colonies were considered for this analysis.

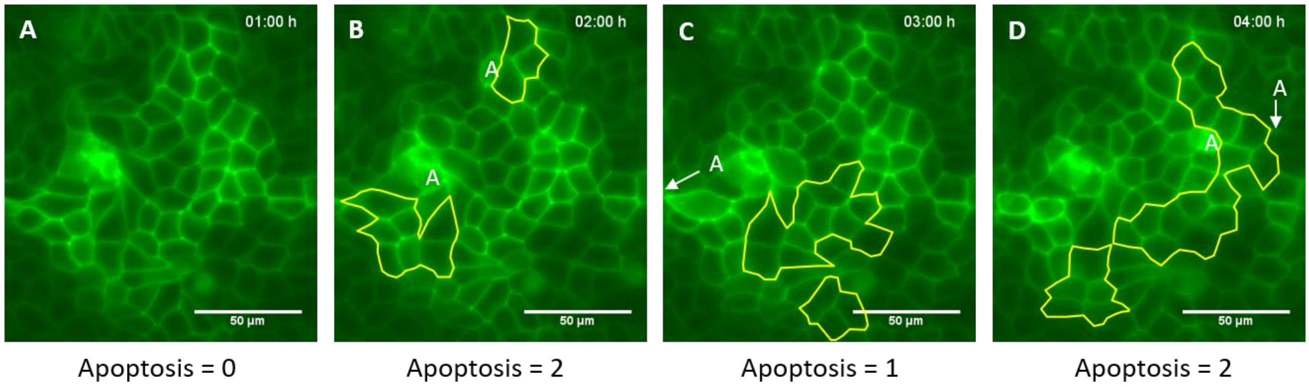


**Figure S8.** Events of apoptosis leads to the emergence of expanding colonies. During (A) 1st hr, (B) 2nd hr, (C), 3rd hr, and (D) 4th hr. Note: The growing cell colonies represented by yellow outline and 'A' represents the location of cells undergone apoptosis.

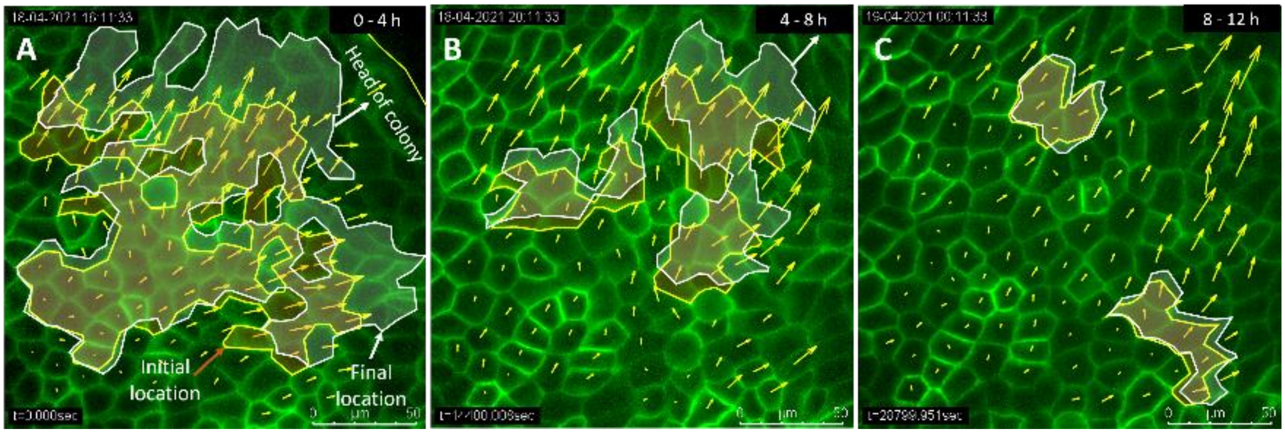


**Figure S9.** The head of Colonies with respect to velocity field vectors (A) 0-4 hr, (B) 4-8 hr, and (C) 8-12 hr. Note: The yellow and white outlines represent the initial and final location of colonies.
